# The BRAIN Initiative Cell Census Network Data Ecosystem: A User’s Guide

**DOI:** 10.1101/2022.10.26.513573

**Authors:** BICCN Data Ecosystem Collaboration, Michael J Hawrylycz, Maryann E Martone, Patrick R Hof, Ed S Lein, Aviv Regev, Giorgio A. A Ascoli, Jan G Bjaalie, Hong-Wei Dong, Satrajit S Ghosh, Jesse Gillis, Ronna Hertzano, David R Haynor, Yongsoo Kim, Yufeng Liu, Jeremy A Miller, Partha P Mitra, Eran Mukamel, David Osumi-Sutherland, Hanchuan Peng, Patrick L Ray, Raymond Sanchez, Alex Ropelewski, Richard H Scheuermann, Shawn Z K Tan, Timothy Tickle, Hagen Tilgner, Merina Varghese, Brock Wester, Owen White, Brian Aevermann, David Allemang, Seth Ament, Thomas L Athey, Pamela M Baker, Cody Baker, Katherine S Baker, Anita Bandrowski, Prajal Bishwakarma, Ambrose Carr, Min Chen, Roni Choudhury, Jonah Cool, Heather Creasy, Florence D'Orazi, Kylee Degatano, Benjamin Dichter, Song-Lin Ding, Tim Dolbeare, Joseph R Ecker, Rongxin Fang, Jean-Christophe Fillion-Robin, Timothy P Fliss, James Gee, Tom Gillespie, Nathan Gouwens, Yaroslav O Halchenko, Nomi Harris, Brian R Herb, Houri Hintiryan, Gregory Hood, Sam Horvath, Dorota Jarecka, Shengdian Jiang, Farzaneh Khajouei, Elizabeth A Kiernan, Huseyin Kir, Lauren Kruse, Changkyu Lee, Boudewijn Lelieveldt, Yang Li, Hanqing Liu, Anup Markuhar, James Mathews, Kaylee L Mathews, Michael I Miller, Tyler Mollenkopf, Shoaib Mufti, Christopher J Mungall, Lydia Ng, Joshua Orvis, Maja A Puchades, Lei Qu, Joseph P Receveur, Bing Ren, Nathan Sjoquist, Brian Staats, Carol L Thompson, Daniel Tward, Cindy T J van Velthoven, Quanxin Wang, Fangming Xie, Hua Xu, Zizhen Yao, Zhixi Yun, Hongkui Zeng, Guo-Qiang Zhang, Yun R Zhang, Jim W Zheng, Brian Zingg

## Abstract

Characterizing cellular diversity at different levels of biological organization across data modalities is a prerequisite to understanding the function of cell types in the brain. Classification of neurons is also required to manipulate cell types in controlled ways, and to understand their variation and vulnerability in brain disorders. The *BRAIN Initiative Cell Census Network (BICCN)* is an integrated network of data generating centers, data archives and data standards developers, with the goal of systematic multimodal brain cell type profiling and characterization. Emphasis of the BICCN is on the whole mouse brain and demonstration of prototypes for human and non-human primate (NHP) brains. Here, we provide a guide to the cellular and spatial approaches employed, and to accessing and using the BICCN data and its extensive resources, including the *BRAIN Cell Data Center (BCDC)* which serves to manage and integrate data across the ecosystem. We illustrate the power of the BICCN data ecosystem through vignettes highlighting several BICCN analysis and visualization tools. Finally, we present emerging standards that have been developed or adopted by the BICCN toward FAIR (Wilkinson et al. 2016a) neuroscience. The combined BICCN ecosystem provides a comprehensive resource for the exploration and analysis of cell types in the brain.

## I. Introduction and Overview

The National Institutes of Health’s Brain Research Through Advancing Innovative Neurotechnologies (BRAIN) Initiative, launched in 2013, is a large-scale effort to accelerate neuroscience research by providing researchers with tools to study and treat human brain disorders through a comprehensive understanding of the human brain (Insel, Landis, and Collins 2013). Following a pilot phase (Ecker et al. 2017), the BRAIN Initiative Cell Census Network (BICCN) launched a major 5-year phase (2017-2022) with the goal of systematic multimodal cell type profiling and characterization of the whole mouse brain, with parallel proof of concept for similar characterization and scalability to tackle the much larger human and non-human primate (NHP) brain. This effort resulted in widespread collaboration among the neuroscience community to apply advanced single-cell profiling to characterize transcriptomic and epigenomic signatures, anatomical phenotypes, and functional properties of brain cell types, and accelerated the normalization of rapid sharing of cell census data with the larger community pre-publication. The success of these efforts built on significant advances in scalable single cell analysis including single-cell genomic (RNA, ATAC-seq and methylation) profiling, anatomical mapping at cellular resolution, and other approaches have proven to be powerful and scalable. BICCN is now on track to deliver on its goal of completing a comprehensive cell census spanning the entire adult mouse brain, and large-scale cell atlas research in human and non-human primate have begun through the new BRAIN Initiative Cell Atlas Network (BICAN), (BRAIN Initiative Cell Atlas Network (BICAN): Comprehensive Center on Human and Non-Human Primate Brain Cell Atlases). The resulting data resources are already proving invaluable for researchers across many areas of neuroscience. Here, we provide a comprehensive description and user guide to these resources and discuss how they can enable rapid progress in neuroscience.

BICCN is a collaborative network of centers and laboratories, including data generating centers, data archives, and data standards developers, which generate, map, and share resources to support several overarching goals. These include generating a high-resolution, comprehensive atlas of cell types in the mouse brain based on large-scale single-cell transcriptome and epigenome sequencing, along with systematic characterization of neuronal morphology, a census of the number and location of cells for each type, new genetic tools to experimentally target brain cell types, and a prototype atlas of human brain and NHP cell types in selected regions of the adult and developing human brain. A standard anatomical template for mapping cell types in the mouse brain was established through completion and validation of a common coordinate framework (CCF, (Wang et al. 2020)). BICCN also conducted an initial profiling of cellular diversity in several structures relevant to neurodegenerative and neuropsychiatric disease including the hippocampus and dorsolateral prefrontal cortex, and importantly, cross-species identification and mapping of cell types between mouse, marmoset, and human (see **Suppl. Materials - BICCN Scientific Outcomes**).

Each BICCN project has contributed publicly accessible data to a multimodal classification of cell types based on transcriptomic, epigenetic, proteomic, morphological connectivity, anatomic distribution, and physiological signatures of cells for further study. To date, the BRAIN Initiative data archives house petabytes of omics, imaging, and neurophysiology data sets generated using over 40 cell profiling techniques and 97 published protocols (see **Sections IV, V**). The BICCN *BRAIN Cell Data Center* (BCDC, biccn.org) manages this ecosystem, together with data archives to support logistical organization, data integration, and development of common data standards as well as central maintenance to sustain, compare and reanalyze data. A major success of the BICCN has been the embrace of a fundamental principle that data should be released on a quarterly basis, pre-publication, and all data is freely shared under CC-BY-4.0 license unless human protection restrictions apply. In this way the BICCN Data Ecosystem represents one of the largest resources for single cell data of the brain and any organ.

The first phase of the BICCN generated a multimodal cell census and atlas of the mammalian primary motor cortex (MOp or M1) (BRAIN Initiative Cell Census Network (BICCN) 2021). This project involved coordinated large-scale analyses of single-cell transcriptomes (Yao, Liu, et al. 2021; Bakken et al. 2021), chromatin accessibility (Y. E. Li et al. 2021), DNA methylomes (Liu et al. 2021), spatially resolved single-cell transcriptomes (M. Zhang et al. 2021), anatomic characterization with morphological and electrophysiological properties (Scala et al. 2020; Muñoz-Castañeda et al. 2021), and cellular resolution input-output mapping (Y. E. Li et al. 2021; Z. Zhang et al. 2021). These results and their extension to the whole mouse brain and other human regions represent a milestone in the effort to create a catalog or census of all brain cell types and advance the collective knowledge and understanding of brain cell type organization. Six active BICCN Working Groups continue to extend and integrate new and existing data across labs towards an integrated transcriptomic and epigenomic atlas of the entire mouse central nervous system.

BICCN reflects the increasingly collaborative nature of modern neuroscience and has accomplished the deepest coordinated characterization of cell types in any organ to date. Consortia such as the Human Cell Atlas (HCA, (Regev et al. 2017)) and Human Biomolecular Atlas Program (HuBMAP Consortium 2019) are strong representatives of this community leading molecular profiling in other organs. Here, we describe the BICCN data ecosystem and provide a guide to accessing and using its data and resources. **Section II** describes the challenge of brain cell type profiling, the approaches taken by BICCN investigators, and requirements for spatial localization and data architecture. **Section III** overviews the data ecosystem and the BCDC and its role in data management. **Section IV** provides a guide to the primary BICCN related data archives describing methods for accessing archived data, and the process of data submission. **Section V** describes progress in standardizing molecular and image-based processing pipelines and their use. In **Section VI** we provide usage vignettes and describe some of the many BICCN tools for analysis and visualization that have been developed. Standards that have been developed or adopted throughout the BICCN are described in **Section VII** and inventory of progress in FAIR (Wilkinson et al. 2016b) neuroscience. Finally, a set of supplementary tables provides information throughout the guide on the many resources available to users of these rich data.

## II. Characterizing Cell Types of the Brain

Understanding reproducible features of brain cells is a prerequisite to characterizing cell types to understanding their function in the brain, to manipulate them in controlled ways, and to understand variability in brain disorders. Neurons can be distinguished by types expressing neurotransmitters, electrophysiological firing patterns, morphology, connectivity, as well as other patterns of gene expression, and these form a natural basis for classification. Properties of glial cells, vascular cells, and immune cells in the nervous system are also essential to understand brain function in health and disease (Sontheimer 2021). Moreover, the brain has an immensely complex global and regional structure and mapping the distribution of cell types across regions and nuclei is a vital part of characterization.

### II.1 BICCN Cell Type Profiling

There is general agreement that types should be defined by invariant and generally intrinsic properties, and that this classification can provide a good starting point for a census (BRAIN Initiative Cell Census Network (BICCN) 2021). There are however significant challenges in characterizing cell types due to inherent biological variability, imperfect measurements, and challenges of data integration between modalities (Mukamel and Ngai 2019). While past attempts have not resulted in a unified taxonomy of neuronal or glial cell types, partly due to limited data, single-cell transcriptomics is enabling, for the first time, systematic high-throughput measurements of brain cells and generation of datasets that hold the promise of being complete, accurate and permanent (Yuste et al. 2020). However, the structure and relationships of cell types is highly complex with evidence indicating that there are not always sharp boundaries separating different regions, particularly in the cortex (Yao, van Velthoven, et al. 2021), (Yao, van Velthoven, et al. 2021; Hawrylycz et al. 2012). For a recent overview of brain cell type profiling and its challenges see (Zeng 2022).

A full characterization of cell types for a given brain region would consist of an enumeration of distinct types characterized across different biological features, including the distributions of their molecular profiles (transcriptome, proteome, chromatin accessibility), developmental history, morphology, functionality (e.g., electrophysiology), and spatial location mapped to a common coordinate framework (CCF) or atlas of the brain. The result of profiling is the development of a taxonomy of different types derived from multiple modality data and their congruences (**Figure 1**). Present classification studies approach but generally do not fully attain complete characterization, having partial measurements and lacking full data correspondences. Determining relative significance of data is challenging and while each modality is valuable, the transcriptome forms a natural template to which other modalities can be mapped for completion and essential missing information.

**Figure 1.**
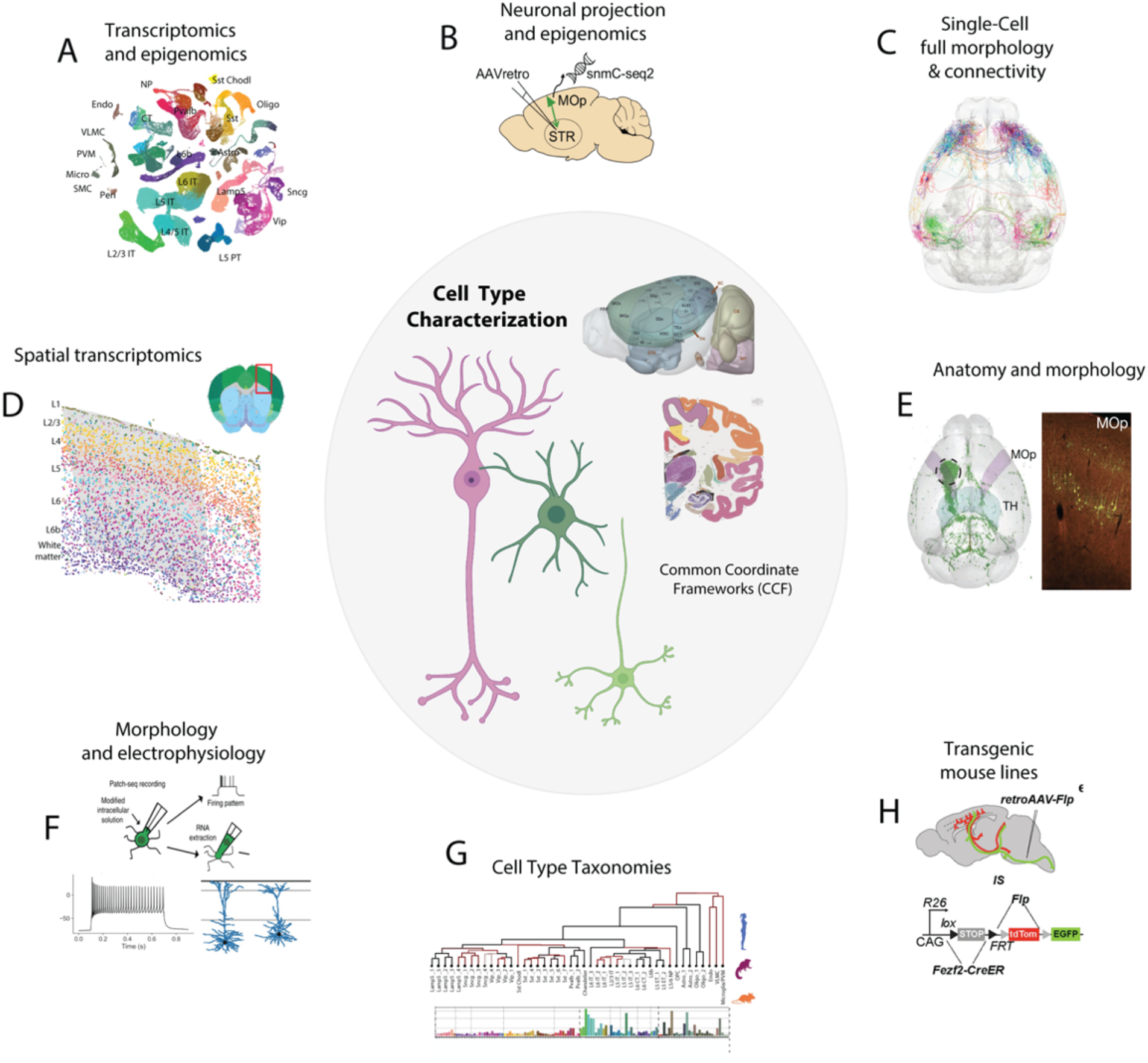
Cell type profiling and major approaches. A variety of multimodal techniques are used to profile cell types of the brain. A Common Coordinate Framework (CCF) is used to map spatial distribution of types and their connectivity. Top to bottom: A) Transcriptomic techniques, single cell and nucleus (sc/sn-RNA-seq), and epigenomic (ATAC-seq), single nucleus methylation (snmC-seq), B) Epi-retro-seq, C) Single cell full morphology and connectivity, (fMOST, BAR-seq), D) spatial transcriptomics (MERFISH), E) antero- and retro-grade tracing methods for morphological reconstruction. F) Multimodal technique combining transcriptome, electrophysiology, and morphology (Patch-seq), H) use of genetic lines. G) Cell type classifications are represented as taxonomies reflecting hierarchical relationships, multimodal correspondence, and cell distribution (**Suppl. Table 1**).

To progress towards this goal, BICCN studies used a wide array of approaches (**Figures 1** and **2**; **Suppl. Table 1**), broadly classified as single cell transcriptomic and epigenomics, spatial transcriptomics, anatomy/morphology, imaging based, electrophysiology, and multimodal, spanning more than 40 high-resolution methods for investigation of cell type characteristics. Some of the most broadly used BICCN methods (**Figure 1**) include single cell and single nucleus RNA-seq (sc/snRNA-seq) (Yao, Liu, et al. 2021; Di Bella et al. 2021; Kozareva et al. 2021; Hodge et al. 2019), single-nucleus long-read sequencing (Hardwick et al. 2022), single cell ATAC-seq, (Yao, Liu, et al. 2021), snmC-seq (Liu et al. 2021), epi-retro-seq (Z. Zhang et al. 2021), single cell full morphology and BAR-seq (Peng et al. 2021), MERFISH and other spatial transcriptomics methods (M. Zhang et al. 2021), anterograde and retrograde tracing for morphology (Muñoz-Castañeda et al. 2021), multimodality Patch-Seq (M. Zhang et al. 2021; Scala et al. 2020), and the use of transgenic lines (Matho et al. 2021). All data in the mouse was mapped to the CCF either through image registration or specimen pinning (**Section II.3**).

The cell type profiling techniques developed and used by the more than 30 BICCN projects are presented in **Figure 2**, illustrating the breadth of the consortium’s approaches. These techniques are broadly classified as transcriptomic, epigenomic, spatial transcriptome, anatomy/morphology, imaging based, electrophysiology, and multimodal, spanning a wide range of more than 40 high-resolution methods for investigation of cell type characteristics. Investigators have been grouped by techniques common to their programs. While the primary focus of the BICCN is on the mouse, **Figure 2** shows profiling applied to human, marmoset, and macaque as well as several other species for an evolutionary study (Bakken et al. 2021). **Suppl. Table 1** provides details on the primary techniques used and BICCN investigator projects (see also **Team Pages** on biccn.org).

**Figure 2.**
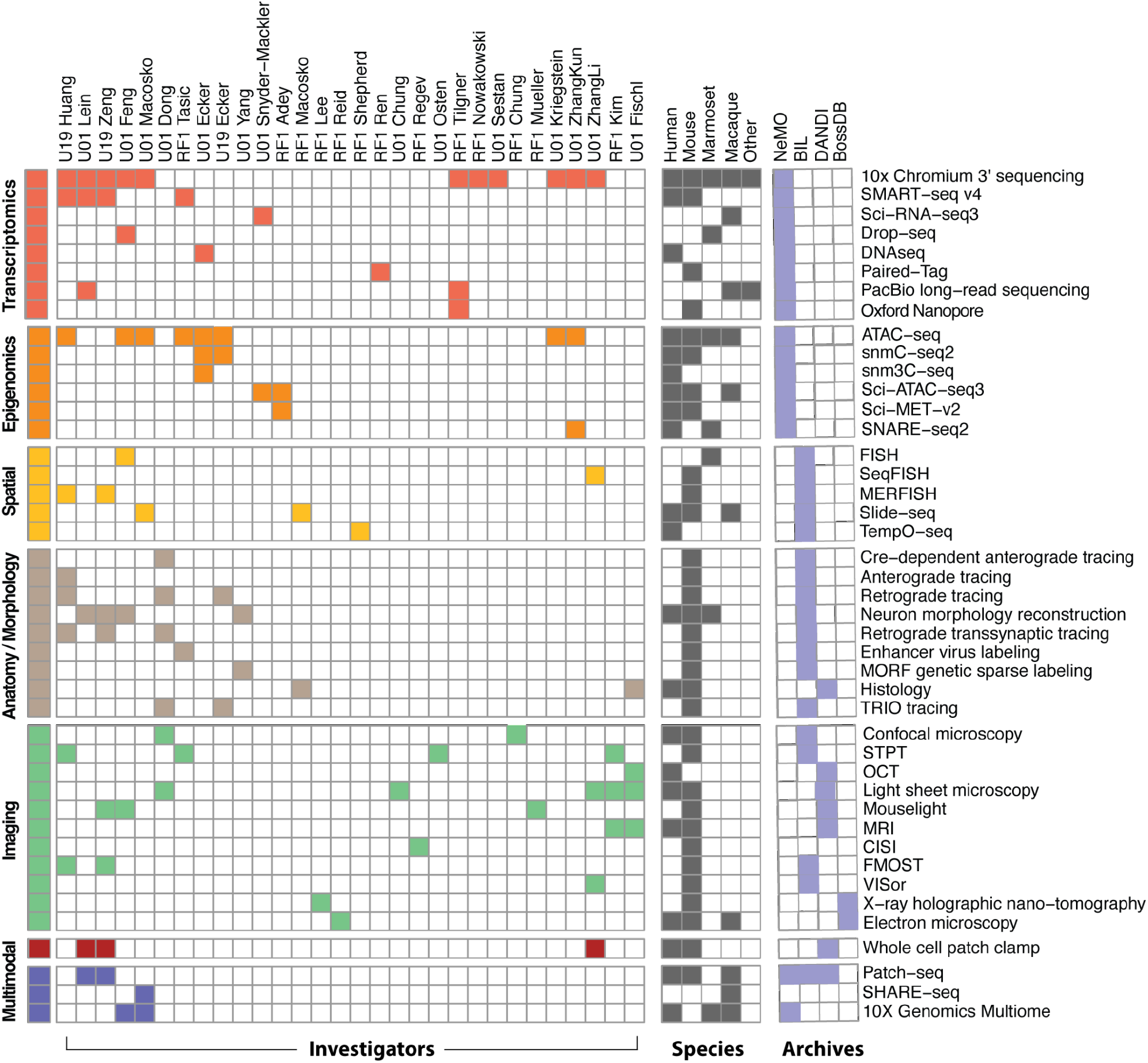
BICCN cell type modalities, techniques, and investigators. Primary techniques (right annotation) used in profiling cell types by BICCN investigators (top) are colored by major modality (left) and primary species (**Suppl. Table 1**). Investigator awards are ordered by techniques common to laboratories. BRAIN Initiative data archives house primary data shown by modality, NeMO: Neuroscience Multi-Omic Archive, BIL: Brain Imaging Library, DANDI: Distributed Archives for Neurophysiology Data Integration, BossDB: Brain Observatory Storage Service and Database (see **Sectio**n IV.) The NIH **UM1**: Cooperative agreements involving large-scale research activities, **U19** multidisciplinary with specific major objective, **U01:** discrete, specified, circumscribed project, **RF1**: discrete, specific project by named investigator (NIH Grants).

### II.2 Data Levels

The importance of having structure in data grows with increasing annotation and association with existing knowledge. The hierarchical organization of information is an active area of bioinformatics (Merelli et al. 2014; Singh 2016). Among other benefits, the specification of the structure of a dataset and its relevant metadata provides a mechanism for efficient retrieval of datasets by users. BICCN data and structured data sets are classified through **Data Levels** (**Suppl. Fig. 1**) reflecting a common conceptual approach for identifying increasing levels of structure from data, through information, to knowledge (Baskarada and Koronios 2013). In this way, BICCN datasets are classified by information content ranging from primary **Raw (Level 0)** data directly from individual laboratories running specific assay platforms, to QC/QA **Validated (Level 1)** data with appropriate associated metadata. **Linked (Level 2)** data that is associated with a specific brain region or nuclei, datasets with computed **Features (Level 3)**, and finally **Integrated (Level 4)** datasets having biological relevant annotation and comparison with other sources (see **Suppl. Materials - BICCN Data Levels**).

Data Levels are more than a classification system and provide an entry point for users of the BICCN data corpus and use-case-directed identification of datasets of particular interest. At project award each BICCN investigator specified levels of data that their project would generate and BICCN working groups collectively reconcile these definitions by each modality such as 10x-snRNA seq, MERFISH, or electrophysiology to achieve modality specific definitions across groups (**Suppl. Table 2)**. While all Level 1 data is required to be deposited in BRAIN Initiative archives (see **Section IV**) on a quarterly basis, uniform storage, and archiving requirements for data sets with more structure are currently being developed by the BRAIN archives as required by BICCN program objectives. **Suppl. Table 3** lists current BICCN level classfied datasets and their provenance. There is flexibility in defining levels particularly with increasing annotation and structure.

### II.3 Common coordinate frameworks of the brain

Spatially localizing cell type data to a CCF provides an anatomic context that is essential to understand the role of cell types in brain function. When mapped in this way, data achieves **Linked** data level, and allows users to identify and access data in a systematic way. BICCN data from the mouse brain is mapped to the *Allen Mouse Common Coordinate Framework* (http://atlas.brain-map.org/, (Wang et al. 2020)), which serves as the main anatomic data browser and spatial coordinate environment for mouse data within BICCN, as well as the reference atlas for mouse data within EBRAINS, the European infrastructure for brain and brain-inspired research (https://ebrains.eu/service/mouse-brain-atlas). CCFv3 is based on a 3-D 10-μm isotropic, highly detailed population average of 1,675 mouse brains using 2-photon imaging (**Figure 3A,B)** and consists of 207 newly drawn structures in 3-D: 123 subcortical structures, 41 fiber tracts (plus ventricular systems), and 43 cortical regions, including primary visual and higher visual areas. Ultimately more than 500 gray matter structures, cortical layers, approximately 80 fiber tracts, and ventricle structures in 3-D will be included. (Wang et al. 2020) (**Figure 3C,D**). A recent fMOST atlas was derived from CCFv3 (Qu et al. 2022)) extending registration accuracy for this modality (**Figure 3E**). The CCFv3 is being refined and improved through the BICCN and currently provides a definitive mouse brain reference framework.

**Figure 3.**
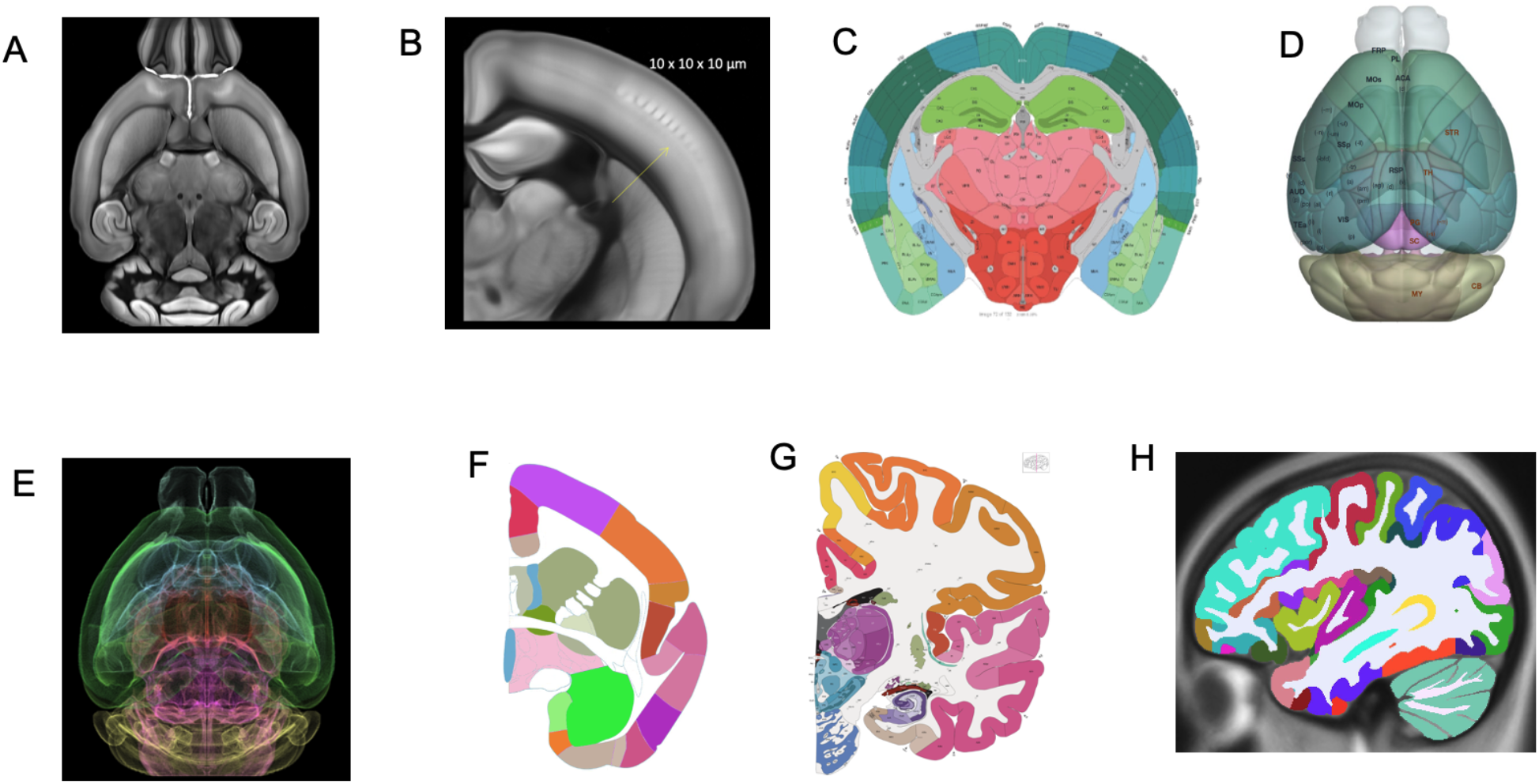
Common coordinate frameworks of the brain. A,B) Allen Mouse Brain Common Coordinate Framework (CCF) constructed from serial two-photon tomography images with 100 μm z-sampling from 1,675 young adult C57BL/6J mice yields 10-*μ*m cubic resolution. C) Digital atlases of the Mouse (Allen CCFv3) annotated plate and D) 3D reconstruction. E) fMOST mouse atlas derived from CCFv3 through iterative averaging of 36 fMOST brains. This approach to a reference atlas reduces the average distance error of somata mapping up to 40% F) marmoset atlas plate (Allen Institute for Brain Science), G) Human reference atlas from 34-year-old female, 1 mm/pixel Nissl and immunohistochemistry anatomical plates, annotated 862 structures, including 117 white matter tracts and several novel cyto- and chemoarchitecturally-defined structures. H) MRI based annotation of human atlas of 150 structures form the initial atlas for BICAN profiling.

Human and non-human primate atlases are similarly necessary for structure identification and data mapping (**Figure 3F-H)**. However, current reference atlases have major limitations such as lack of whole-brain coverage, relatively low image resolution, and sparse structural annotation. The BICCN uses the Allen Human Reference Atlas - 3D, 2020, a human brain atlas (Ding et al. 2017) that incorporates neuroimaging, high-resolution histology, and chemo architecture across a complete adult female brain, with magnetic resonance imaging (MRI), diffusion-weighted imaging (DWI), and 1,356 large-format cellular resolution (1 mm/pixel) Nissl and immunohistochemistry anatomical plates (**Figure 3G)**. It is annotated in 3D to over 150 structures (**Figure 3H**) and has been re-released under a CC-BY-4.0 license in 2022 to support broader community use. This human atlas forms the starting anatomic context for the next BICAN phase.

Mapping brain data to reference spaces is complex and uses a range of manual and automated methods of image registration (see **Section V.2**). Given the challenge of determining anatomical context for any reference atlas even within the mouse, precise image registration requires the whole brain image series (or a reasonable fraction of the brain) to be present with sufficient distinctive anatomical landmarks. Omics data may not have detailed structural localization and can be positioned within a CCF through coordinate based, visual, or ontological tagging. Human data is typically of this type, where anatomic ontology is known and localized using annotated atlas plates and MNI space from the MRI reference brain volume, (ICBM 2009b Nonlinear Symmetric, (Mai and Majtanik 2017). Mapping of these tissues is effectively done using the *Cell Locator* (RRID:SCR_019264), developed in collaboration with Kitware (www.kitware.com). See **Suppl. Materials - Common Coordinate Frameworks** for more information.

## III. The BICCN Data Ecosystem

The BICCN data workflow includes three major components, from work in individual centers, followed by ingestion and storage in dedicated archives, and ending in data catalog and portal in the *BRAIN Cell Data Center* (BCDC) (**Figure 4**). Multimodal data is generated by laboratories in multiple centers, which develop and apply robust methods for high-resolution, high-throughput mapping, including laboratory specific QC/QA methods and data quantification (**Figure 4A**). Data analysis in individual laboratories is focused on rigorous signal detection and clustering, identification of modality specific cell type taxonomies and the validation of cross-modality associations. BICCN mandates broad and rapid data dissemination to accelerate scientific exploration and encourage community engagement, and all laboratories deposit Level 1 validated data quarterly to dedicated archives (**Figure 4B**). Finally, the BICCN data ecosystem is managed by the BCDC (**Figure 4C**). BCDC provides public access to, and organization of the complex data, tools, and knowledge derived by BICCN, by supporting the acquisition of data from BICCN partners, providing data models and framework for importing structured data into the BCDC, and establishing semantic and spatial community standards for description and management of single cell data modalities.

**Figure 4.**
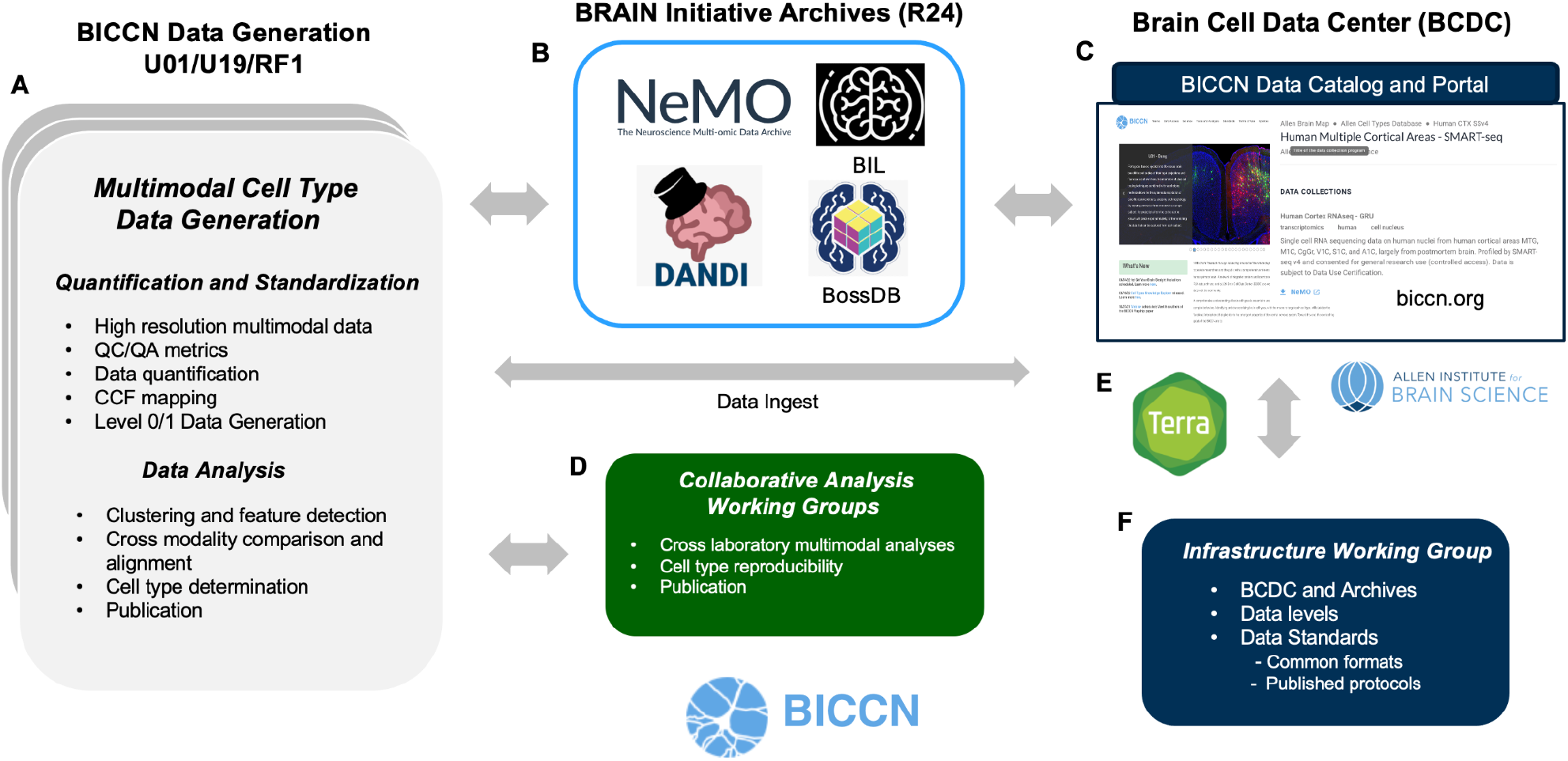
BICCN Data Ecosystem. A) Multimodal cell type data generation by UM1/U01/19, RF1 centers produce high resolution Level 1 multimodal data. B) Data are submitted to one of four BRAIN archives depending on data type(s): Neuroscience Multi-Omic Data Archive (NeMO), Brain Imaging Library (BIL), Distributed Archives for Neurophysiology Data Integration (DANDI) for neurophysiology data, and Brain Observatory Storage Service and Database (BossDB) for electron microscopy ultrastructural datasets. Datasets are indexed and referenced C) by the Brain Cell Data Center (BCDC, biccn.org) which provides a portal for accessing the consortium’s data, tools, and knowledge. D) Laboratories engage in collaborative cross-modality interpretation of data and results. E) Terra cloud-based platform for standardized omics processing accessible through BCDC. F) An infrastructure working group oversees architectural development and workflow management.

BICCN has a highly collaborative network for addressing multimodal analysis and cell type reproducibility across modalities and laboratories (Booeshaghi et al. 2021; BRAIN Initiative Cell Census Network (BICCN) 2021) **(Figure 4D**). Cross institution analysis working groups tackle regional and whole brain analysis, which is facilitated by unrestricted access to quarterly released data to the archives. Systematic data processing provides a platform with common computational pipelines and environment for reproducible science across groups. For example, the BCDC provides access to Terra (https://terra.bio), a scalable and secure platform co-developed by the Broad Institute, Microsoft and Verily for biomedical researchers to share data and run analysis tools such as omics processing pipelines (**Figure 4E**). In addition, an Infrastructure and Standards Development group develops needed software, formalizes cross modality standards, and specifies data structures, and protocols (**Figure 4F**) (see **Section VII**).

The BICCN Portal (www.biccn.org) is an entry point for BICCN resources and provides detailed investigator profiles, consortium news, data access, tools, standards documentation, policies, and overview of scientific progress (**Figure 5A-D**). BCDC maintains a searchable data catalog listing all public datasets available through the BICCN portal. The BICCN catalog is built as an extension of the Allen Institute’s Brain Knowledge Platform and is currently accessible under the “Data access” tab at BICCN.org https://biccn.org/data. The catalog organizes datasets by projects, each with one or more associated datasets that may be stored in a single archive or distributed across multiple archives (see **Section IV**). Users can browse the catalog or use a flexible search function (**Figure 5C**) to filter data by species, modality, techniques, and specimen type. For each dataset, the catalog provides basic descriptive metadata (**Figure 5B**), information on the dataset release status, the terms of the use and a link to the location in the archive. Clicking on the link brings the user to a landing page that provides the dataset identifier, descriptive metadata, a download link, and additional relevant information.

**Figure 5.**
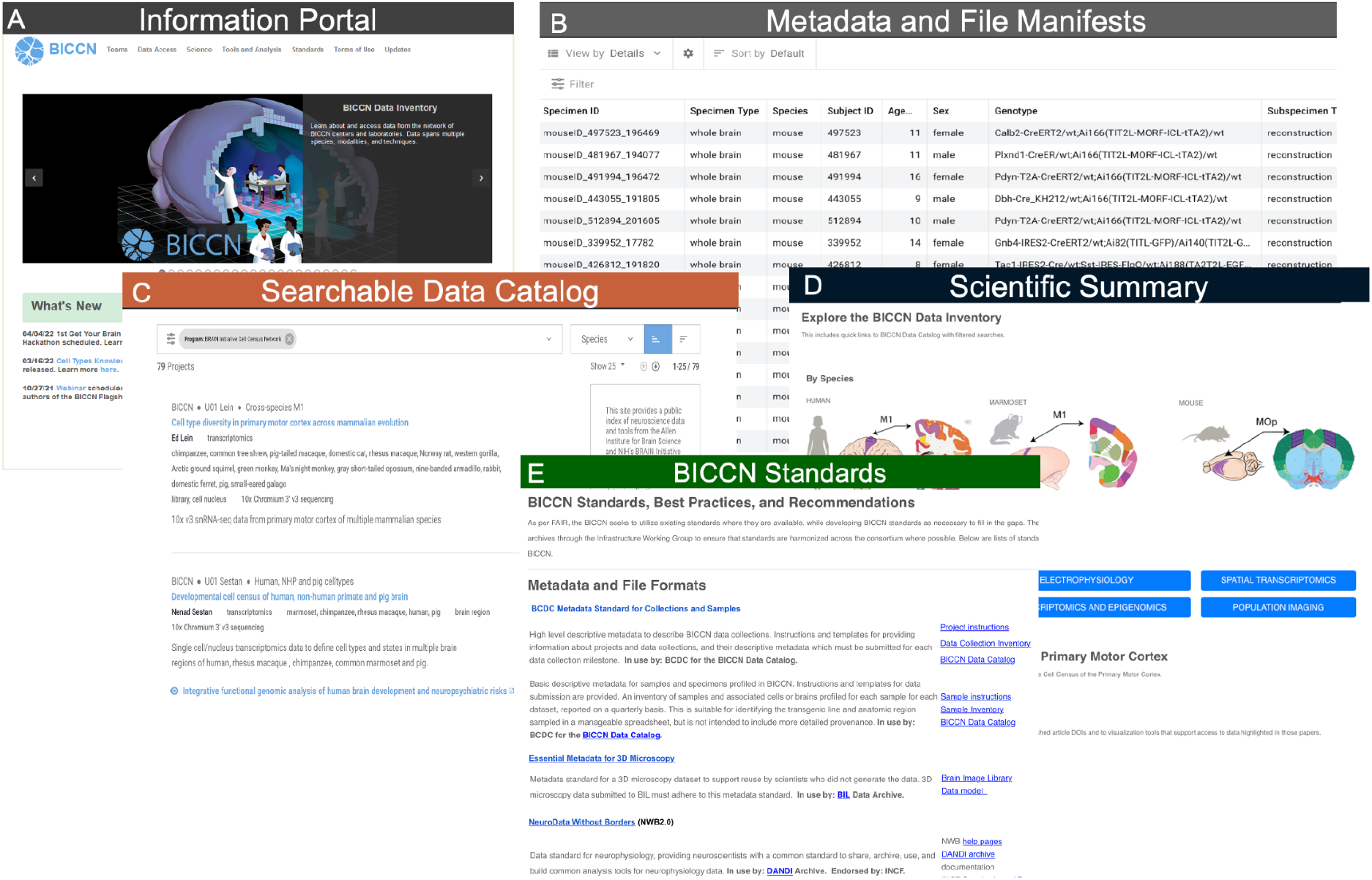
Brain Cell Data Center (BCDC, www.biccn.org). The BCDC supports the goals of the BRAIN Initiative Cell Census Network (BICCN) by providing a central public resource through the BICCN portal A) which makes BICCN data and activities searchable from inside or outside the BICCN network. The portal includes B) Metadata and file manifests documenting data deposition from investigators into archives, C) A searchable data catalog describing projects and datasets generated by the BICCN, D) Links to relevant publications with associated data sets and data mining tools, and E) BICCN standards adopted by the consortium or created by internal working groups to ensure that data are harmonized across the consortium.

## IV. Data Archives for the BICCN

BICCN data archives ensure that data is Findable, Accessible, Interoperable, and Reusable (FAIR) (**Section VII**), while optimizing storage costs to house and process data, enabling reproducible data practices, and effectively managing interchange between data producers and computational analysts. The archives serve as active repositories with adjacent compute capabilities that enable collaboration within and across labs and serve as an entry point for research for all neuroscientists. Each archive supplies its own documentation on data submission, access, and reuse. There are currently four archives that are central to BICCN related data types (out of seven BRAIN Initiative supported archives): Neuroscience Multi-Omic Data Archive (NeMO, https://nemoarchive.org), Brain Imaging Library (BIL, https://www.brainimagelibrary.org), Brain Observatory Storage Service and Database (BossDB, https://bossdb.org), and Distributed Archives for Neurophysiology Data Integration (DANDI, https://dandiarchive.org) (**Figure 6**). The archives supply permanent and archival storage capabilities for transcriptomic and epigenomic data, imaging-based data including tracing, slice and whole brain morphology, density distribution, electrophysiology, and functional imaging, and ultra-resolution electron microscopy (see **Figure 2**).

**Figure 6.**
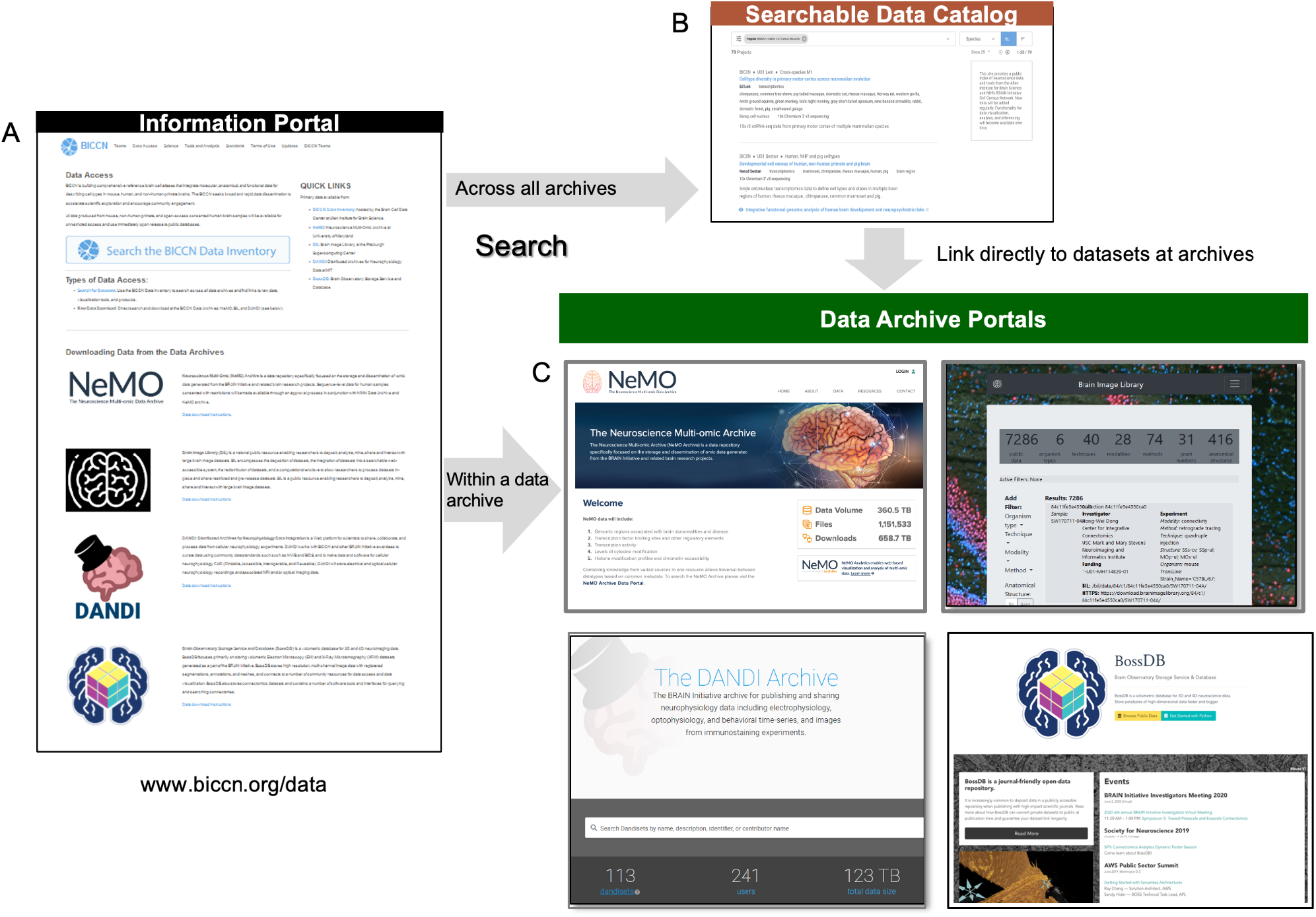
BRAIN Initiative Data Archives and the BICCN. A,B) BICCN information portal provides searchable access to indexed datasets at individual archives. Four primary archives support the BICCN. C) Neuroscience Multi-mic Archive (NeMO) for transcriptomic and epigenomic datasets, Brain Image Library (BIL), Distributed Archives for Neurophysiology Data Integration (DANDI), and Brain Observatory Storage Service and Database (BossDB) (**Suppl. Materials**).

### IV.1 Neuroscience Multi-Omic Archive (NeMO, RRID:SCR_016152, https://nemoarchive.org)

stores and disseminates omics data from the BRAIN Initiative and related brain research projects. NeMO stores both transcriptomic and epigenomic data, including transcription factor binding sites and other regulatory elements, histone modification profiles and chromatin accessibility, levels of cytosine modification, and genomic regions associated with brain abnormalities and disease. Data are organized by projects and within each project further organized by laboratory where data was generated, grant, organism, and the assay type. NeMO is consistent with the principles advanced by the NIH Strategic Plan for Data Science (“NIH Strategic Plan for Data Science” n.d.), including FAIR Principles, documentation of APIs, data-indexing systems, workflow sharing, use of shareable software pipelines and storage on cloud-based systems.

Data archived at NeMO include raw sequence files as well as derived intermediate files such as BAM files (BAM | Integrative Genomics Viewer) and analyzed results including counts and cluster information (**Data Levels 1,3**), and metadata are submitted to BCDC. Sequence-level data for human samples submitted with access restrictions are made available through an approval process in conjunction with the NIMH Data Archive and NeMO archive. Data submissions rely on the fast transfer technology Aspera (https://www.nemoarchive.org/resources/aspera/). To upload data to NeMO, a user obtains credentials through an account request form (https://nemoarchive.org/register.php). Submitters of restricted access data use account usernames with an appended “-restricted”. All submissions begin with the upload of a tab delimited (.tsv) file manifest, including MD5 checksums through the User Dashboard at https://nemoarchive.org/login. Submitted data are placed in an “incoming” area, while they are reviewed for quality control. The NeMO Data Portal provides a simple filtered search interface to help identify data of interest. The BICCN portal (biccn.org) provides direct access to NeMO and data can also be downloaded from the NeMO website using HTTP.BICCN Omics Workshops have provided a good source for instructional tutorials (“BICCN Omics Workshop 2022” n.d., “BICCN Omics Workshop 2022” n.d.).

### IV.2 Brain Image Library (BIL, *RRID*: SCR_017272, https://www.brainimagelibrary.org)

provides a persistent centralized repository for brain microscopy data, and supports dataset deposition, integration into a searchable web-accessible system, redistribution, and analysis tools. It allows researchers to process datasets in-place and to share restricted and pre-release datasets. BIL includes whole-brain microscopy image datasets and their accompanying higher level derived data such as neuron morphologies, targeted microscope-enabled experiments including connectivity between cells and spatial transcriptomics, and other historical collections of value to the community. More generally than the BICCN. BIL accepts all microscopy data relevant to the BRAIN Initiative, including data from primates, most mammals and model organisms. BIL accepts both raw and processed data, and Data Levels 2-3, such as neuron tracings, can be linked to lower-level data sources. While BIL does not limit the amount of data deposited per dataset or investigator, users planning to deposit more than 50TB of data in a single year should contact in advance to discuss data deposition plans. Data contributed to BIL following the Standard metadata for 3D microscopy schema (Ropelewski et al. 2022) are issued DOIs. Higher-level traced neuron data is accepted in the SWC format (“NeuroMorpho.Org - a Centrally Curated Inventory of Digitally Reconstructed Neurons and Glia” n.d.).

The BIL Analysis Ecosystem provides an integrated computational and visualization system to explore, visualize, and access BIL data without having to download it. The Analysis Ecosystem provides large memory nodes, GPU nodes, and access to high-performance computing (HPC) resources for extensive data exploration. The Analysis Ecosystem virtual machine (VM) system has a remote desktop environment to run applications such as Fiji (Schindelin et al. 2012) and Vaa3d (Peng et al. 2010), and supports custom web gateways and commercial software. An Open-OnDemand gateway at BIL offers interactive access to popular scientific applications such as Jupyter Notebooks. A search portal (**Figure 6C**) provides pointers to the data on the BIL Analysis Ecosystem as well as download links. Finally, workshops are offered on a regular basis on how to interact with data through the BIL Analysis Ecosystem, the data submission process, and additional services (“Brain Image Library: Contact” n.d., “Brain Image Library: Contact” n.d.).

### IV.3 Distributed Archives for Neurophysiology Data Integration (DANDI, *RRID*:SCR_017571, https://dandiarchive.org)

is a Web platform for scientists to share, collaborate, and process data from cellular neurophysiology experiments. DANDI works with BICCN and other BRAIN Initiative groups to curate data using community data standards such as Neurodata without Borders (NWB, (Rübel et al. 2022) and Brain Imaging Data Structure (BIDS, (Gorgolewski et al. 2016)), and to make data and software for cellular neurophysiology FAIR. Currently housing over 400 TB of data across 6 species and multiple instruments and techniques, the DANDI archive stores, publishes, and disseminates neurophysiology data including electrophysiology, optical physiology, and behavioral time-series, and images (MRI and microscopy) from immunostaining experiments.

DANDI datasets are referred to as *Dandisets* and include dataset and file metadata. Supplied per-file metadata includes instrument, species, sample, subject, and other experimental details. Each Dandiset is organized in a structured manner to help users and software tools interact with it, and has a unique persistent identifier which can be used for citation. The DANDI Web application allows users to browse and search for Dandisets, create an account to register a new Dandiset or gain access to the Dandi Hub analysis platform, add collaborators to a Dandiset, and retrieve an API key to perform data upload. DANDI has enabled streaming access to parts of data using a combination of cloud technologies and storage formats, allowing for more scalable analysis software and visualization technologies. DANDI exposes all data as versioned DataLad datasets (Halchenko et al. 2021), allowing users to overview an entire dataset without downloading any data to their local filesystem, and then to selectively download specific files or folders. DANDI provides a programmable interface to the archive and Jupyter computational environment, and an API allows development of other software tools for accessing, searching, and interacting with the data in the archive.

## V. BICCN Data processing pipelines

### V.1 BICCN molecular pipelines

The BICCN ecosystem includes production-level, cloud-native data processing pipelines, developed by the Broad Institute’s Data Sciences Platform (DSP) in collaboration with BICCN. While BICCN investigators and other users often process their own omics datasets, standardized pipelines are used to supplement and integrate original analyses with uniformly processed data sets. The pipelines leverage consistent standard file schema and types as well as standardized quality control metrics and metadata. The established cloud-native pipelines replicate processing used by several consortium groups including computational pipelines for processing single cell/nucleus 10x v2/3, sc/sn full transcript, sn-ATAC-seq, and snmC-seq sequencing data. Each of these pipelines was developed in collaboration with a sponsoring BICCN group and captures their expertise in data processing (**Table 1**; for additional pipeline documentation, type “BICCN” in the **WARP** Documentation search bar (https://broadinstitute.github.io/warp/docs/get-started).

**Table 1.**
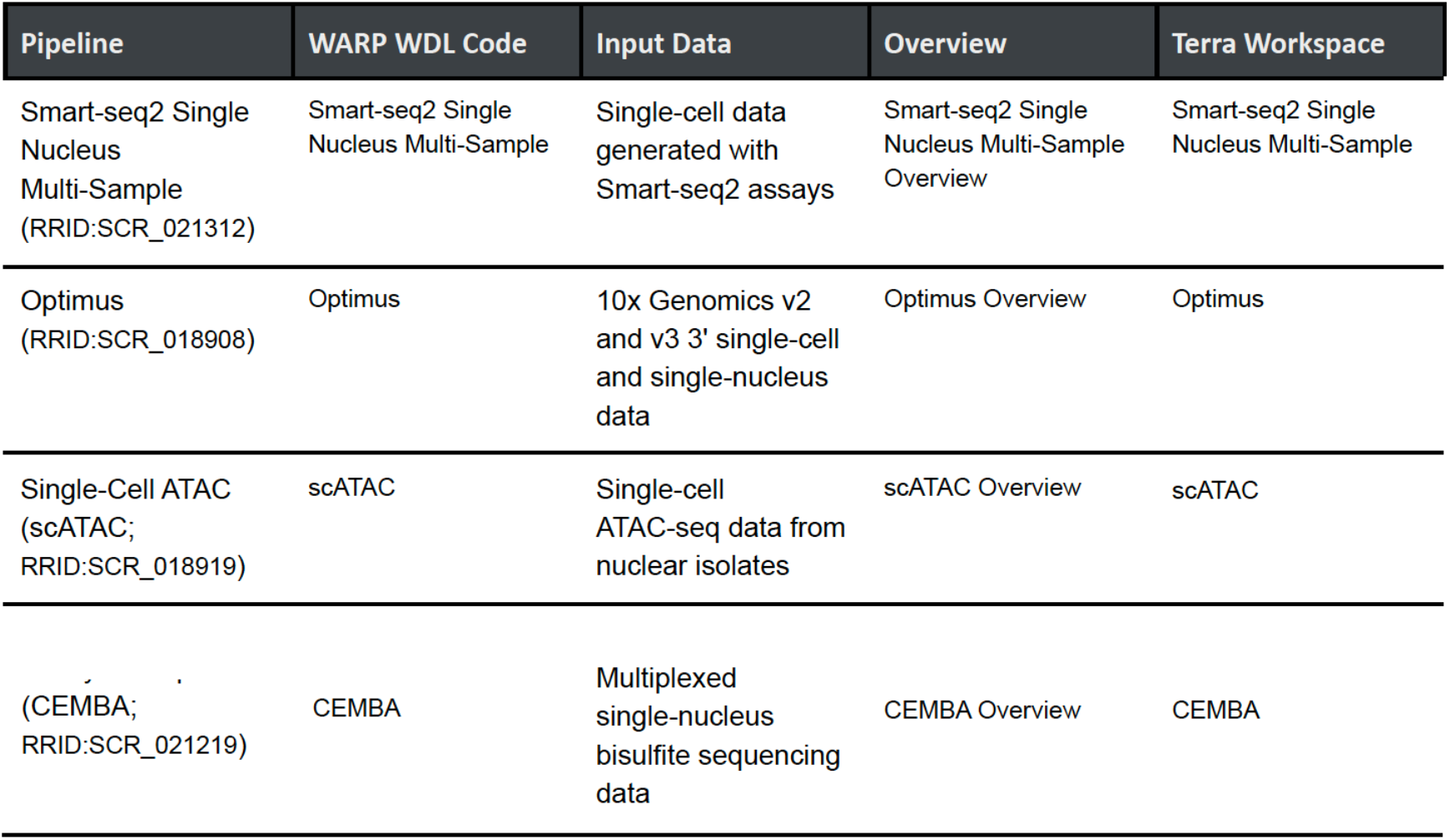
BICCN Molecular Pipelines. A systematically developed set of pipelines for processing molecular omics data is under continuous development by the Broad Institute and BICCN team. Computational pipelines are available for processing single cell/nucleus (sc/sn) 10x v2/3 3 prime, sc/sn full transcript, sn ATAC-seq, and snmC-seq sequencing. Detailed documentation and user guides are available through supplied links. Workflow Description Language (WDL) Analysis Research Pipelines (WARP WDL Code) repository contains a collection of cloud-optimized pipelines.

The Broad Institute Data Sciences Platform resources are actively used by other individuals and consortia, and the approach to the development of molecular pipelines for the BICCN is inspired by FAIR principles (Wilkinson et al. 2016b). This includes use of Research Resource Identifiers (RRIDs) to give pipelines unique, explicit identifiers, and hosting the pipelines in multiple community resources including: public GitHub repositories (for software engineers), Dockstore (https://dockstore.org) for computational biologists, and Terra (where the pipelines are pre-configured and ready to run for those without local infrastructure or who want to use scalable cloud resources). Use of a modern Workflow Description Language (WDL) separates the code performing scientific tasks from code orchestrating the pipeline on infrastructure and encouraging interoperability for reproducible science (see **Section VII**).

### V.2 BICCN Image processing pipelines

Image registration, mapping, and alignment are necessary to bring data from individual brain samples into common coordinate systems, yet often challenging to standardize. The choice of which software to use often depends on the computational resources available, integration with other image tools (e.g. visualization, neuron reconstruction), and which algorithm is most effective for the image data at hand. Two major image processing platforms were developed or extended through the BICCN data ecosystem. Generative Diffeomorphic Mapping (GDM) for image registration and atlas mapping from the Brain Architecture Portal (http://brainarchitecture.org) combines multimodal imaging datasets such as *ex vivo* radiology and histology in the same animal/subject). The GDM approach overcomes challenges in registering tissue processing procedures such as extraction and fixation that cause brain tissue deformation (**Suppl. Materials - CCF Mapping**). The Image and Multi-Morphology Pipeline (Y. Li et al. 2022; Peng et al. 2010) accesses raw images from the Brain Image Library archive (**Section IV.2**) and implements the full pipeline of conversion, processing, morphometry generation, registration and mapping, release, and analysis. The pipeline is hosted on an open cloud platform that features collaborative processing and synergetic computing among various clients, and web interfaces. All data on the server can be accessed through MorphoHub (Jiang et al. 2022), a petabyte-scale multi-morphometry management system and integrates the three largest whole-brain full morphology datasets (Peng et al. 2021), MouseLight (Winnubst et al. 2019), and single-neuron projecome of mouse prefrontal cortex (Winnubst et al. 2019; Gao et al. 2022) (**Suppl. Materials - Image Processing Pipelines**).

While not all BICCN image data is mapped into the CCFv3, a wide variety of tools were improved through BICCN collaboration **(Suppl. Fig. 2**). These image registration packages include ANTs, which maximizes image cross-correlation while ensuring that maps between images are smooth and invertible (Avants et al. 2008), Elastix, whose modular design allows users to compare different registration algorithms (Klein et al. 2010), and 3D Slicer, which offers both landmark and grayscale image based registration (Fedorov et al. 2012). Highly flexible registration tools such as QuickNII and VisuAlign directly map to the CCFv3 and focus on registering high resolution 2D images (Puchades et al. 2019), CloudReg which is a cloud-compliant pipeline for intensity correction, image stitching, and diffeomorphism based registration (Chandrashekhar et al. 2021), and mBrainAligner (Y. Li et al. 2022), which is cross-modal, and integrates with the Vaa3D software suite and can also be freely accessed through the web server mBrainAligner-Web (Y. Li et al. 2022; Qu et al. 2022). Additional cloud-based Petabyte data generation and management system MorphoHub (Jiang et al. 2022) was also developed to assist additional data analysis. These platforms are developed through open-source and extensible approaches, are accessible to the public, and can be extended through plugins (**Suppl. Table 4**).

## VI. Working with BICCN Data

The BICCN has developed many tools and applications to work with BICCN data. An inventory of these tools describing their application to single cell analysis is provided under the “Tools and analysis” tab of the BICCN portal Tools and Analysis - Brain Cell Data Center (BCDC). Some of these resources are described below, with an emphasis on those that facilitate integrative analysis.

### VI.1 Cell Type Knowledge Explorer

The *Cell Type Knowledge Explorer (CTKE*, RRID:SCR_022793) is an interactive application that aggregates multimodal BICCN data from the primary motor cortex (MOp) atlas at the level of individual cell types in mouse, human and marmoset. The CTKE integrates the work of many BICCN laboratories and presents aggregate knowledge about cell types in the form of data visualizations and text summaries. Drawing inspiration from Gene Cards (genecards.org (Safran et al. 2010)), this information is displayed on over 400 individual panels across the three species. The CTKE is powered by a data-driven ontology (Tan et al., n.d.) linking MOp atlas data to a well-established body of knowledge on neurobiology enabling text-based search of the data by cell type names, minimal sets of marker genes from the NS-Forest algorithm (Aevermann et al. 2021), and historical terms from the literature (e.g., “pyramidal”, “chandelier”, etc.).

By leveraging BICCN’s cross-modality mapping of non-transcriptomic data to expression-based taxonomies, CTKE provides rich phenotypic information about cell types and enables its systematic exploration. *Cell Type Knowledge Cards* for each of the three taxonomies are accessible from an interactive sunburst plot (**Figure 7A,B**). Each card visually presents the molecular signatures of cell types derived from single-cell transcriptomics, and may also include morphological reconstructions, exemplar action potential traces and summaries of electrophysiological characteristics; or spatial locations determined using spatial transcriptomics. Genome browser views show accessible chromatin data at marker gene locations and predicted cell type-specific enhancer regions and links to homologous cell types across species. These modalities are represented on any given card in unique panels, each of which includes links to reusable source data and additional BICCN visualization and analysis tools.

**Figure 7.**
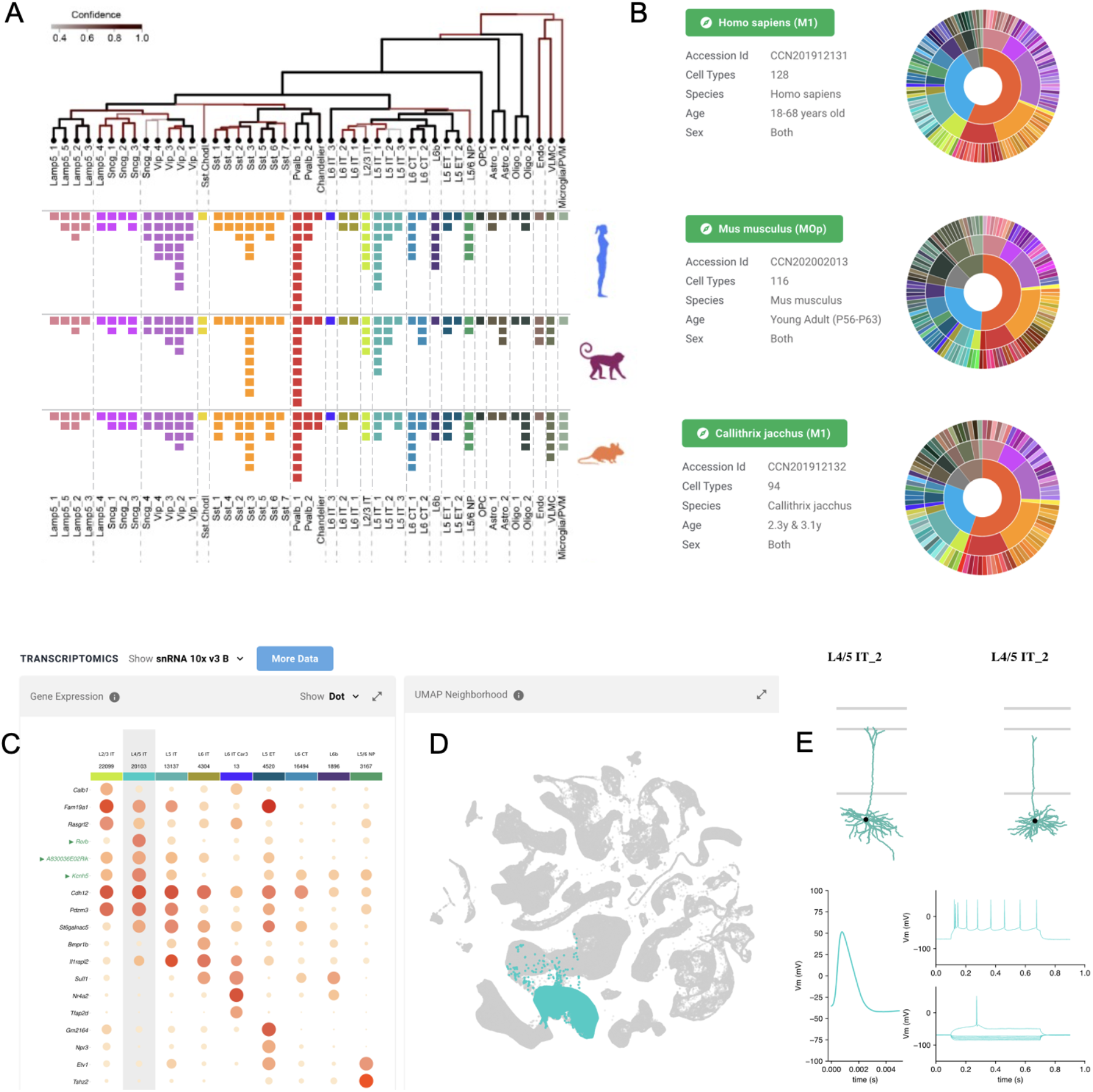
Cell Type Knowledge Explorer. CTKE is an interactive tool that aggregates multimodal BICCN data from the primary motor cortex mini atlas at the level of individual cell types in mouse, human and marmoset. A) Cross-species aligned taxonomies and common cell types in MOp, B) Cell types in each species are accessible and linked through interactive sunburst plots, C) Marker gene panels defining the L4 cell type in mouse, including NS-Forest markers (Aevermann B., et al. 2021) highlighted in green, and D) rendering of expressing cells in UMAP, E) Morphology and electrophysiology exemplars associated with cell type.

CTKE also helps researchers to annotate and interpret their own data. For example, CTKE includes links to Azimuth (Hao et al. 2021), a web application that provides utilities to map single-cell expression data to curated reference datasets. This allows users to derive cell type annotations for their own datasets in the context of BICCN primary motor cortex mini atlas data for the human and mouse. Similarly, CTKE facilitates the interpretation and annotation of other data types. For example, a researcher studying mouse MOp may have immunohistochemistry data indicating that the gene *Rorb* is highly expressed in a certain population of cells and want more information about what type of cells they might be. Searching “*Rorb”* in the CTKE would return the L4/5 IT neuron subclass as a cell type that expresses *Rorb* more highly than other MOp types (**Figure 7C,D**, “Transcriptomics” panel). Navigating to the “Spatial Transcriptomics” and “Morphology” panels might reveal that L4/5 IT neurons are found at a similar cortical depth and with similar morphological characteristics to those this researcher sees in their cell population of interest (**Figure 7E**). If this researcher were interested in understanding whether these cell types are present in humans, they could navigate to the “Cross-Species Cell Types” panel on the Cell Type Knowledge Card for the L4/5 IT_1 subtype, where they would also find several putatively homologous types and be able to navigate directly to their cards for further investigation. In summary, the CTKE strives to provide a user-friendly interface for deep exploration of the BICCN primary motor cortex mini-atlas in a cell type-centric manner and provides a framework for extending to other brain regions and future data navigation tools for whole-brain multimodal atlases.

### VI.2 NeMO Analytics (RRID:SCR_018164; **https://nemoanalytics.org)**

is a web-based suite of data visualization and analysis tools for single-cell data analysis. The portal allows users to explore single-cell, single-nucleus and spatial transcriptomic and epigenetic profiling data, with flexible plotting tools allowing side-by-side comparisons of any data type. The portal is pre-populated with thematically organized datasets reflecting projects across BICCN. Users can upload their own data for private or public use, utilize curated datasets from other users, select a dataset from the NeMO Archive, or benefit from data collections hosted from peer-reviewed publications. NeMO Analytics simplifies access to BICCN data and provides non-programmers with a suite of analytical tools for data exploration, including: cell cluster visualization based on expression/cell type, cell cluster comparison, identification of marker genes across datasets, plotting multi-gene analyses (e.g., heat maps, volcano plots, violin plots for groups of genes), and note taking (**Figure 8**). Additional tools include a workbench to perform *de novo* analysis of scRNA-seq data, visualization and analysis of spatial transcriptomics data, and visualization of epigenomic data. The platform supports visualization of datasets across species and modalities side-by-side and linked by homologous gene symbols. The links to the datasets in NeMO Analytics are embedded in the figure legends, providing seamless access to the data.

**Figure 8.**
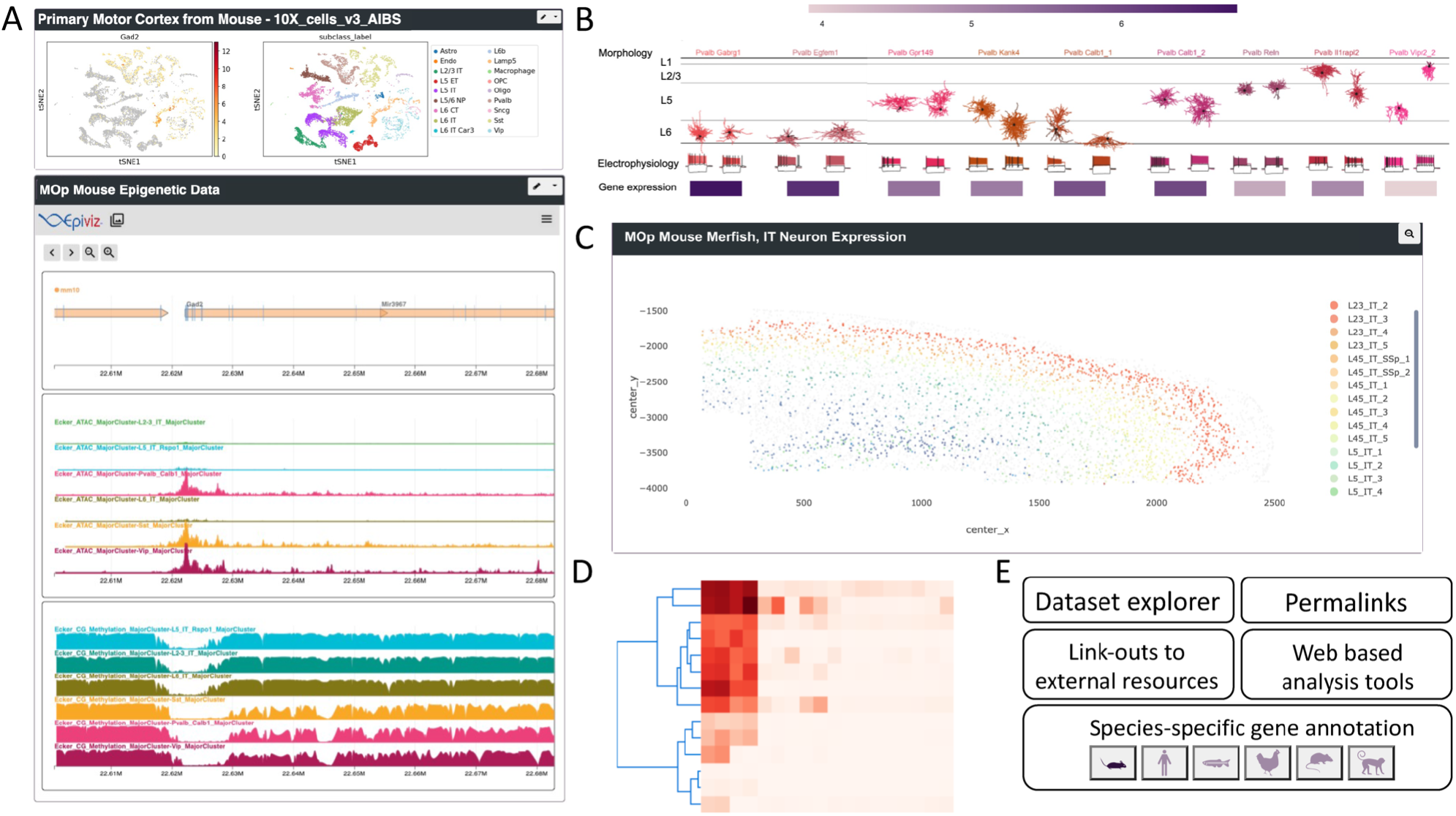
Analyzing BICCN datasets with NeMO Analytics. NeMO Analytics provides direct access to many of the BICCN multi-omic datasets for comparative analysis, visualization and data mining. A) Example NeMO Analytics profile showing glutamic acid decarboxylase 2 (*Gad2*) expression and epigenetic changes in the datasets of (Yao, Liu, et al. 2021). This profile can be found at NeMO Analytics B) Integrated visualization of Patch-seq morphology, electrophysiology, and gene expression for cell types in primary motor cortex NeMO Analytics (Scala et al. 2020). C) Visualization from a MERFISH experiment with spatial distribution of cell types NeMO Analytics (M. Zhang et al. 2021). D) NeMO Analytics offers a variety of web-based visualization and analysis tools including heatmaps, volcano plots, and a single cell workbench allowing for *de novo* analysis of datasets. E) Additional utilities of NeMO Analytics.

### VI.3 Mouse Connectome Project (MCP, *RRID*:SCR_004096, https://cic.ini.usc.edu; https://brain.neurobio.ucla.edu/)

The MCP has systematically produced and collected connectivity data for over 10,000 neural pathways in >4000 experimental cases utilizing a variety of multi-fluorescent pathway tracing techniques that included double coinjections, triple anterograde and quadruple retrograde tracing, Cre-dependent double AAV anterograde tracing, and rabies viral based Cre-dependent retrograde methods (Bienkowski et al. 2018; Zingg et al. 2014), (Benavidez et al. 2021a), (Foster et al. 2021) (Figure 9). This combination of injection strategies can (Benavidez et al. 2021b) simultaneously reveal key connectivity information for a given brain region and enables construction of detailed connectivity maps and to systematically assemble neural networks of different functional systems in the mouse brain. Complementary to the molecular cell typing strategies described above, these connectivity data provide a fundamental framework for cataloging neuronal types based on anatomic locations, projection targets, and morphological features (**Fig. 9A**). In each animal, up to four retrograde tracers are injected into different cortical locations to retrogradely label all neurons that send projections to the injected areas. Because the injections collectively span the entire neocortex, theoretically, for any given cortical area, all neuronal populations (corticocortical projection neurons) that innervate different cortical targets have been demonstrated. Distributions of these retrogradely labeled neurons were annotated to construct a connectivity matrix to visualize corticocortical network organization and a connectivity map to enable direct comparisons of regional and laminar specific distribution patterns of neuronal populations (cell types) associated with each cortical area (**Fig. 9C**); see also www.MouseConnectome.org/Corticalmap). Multiple retrograde tracer injections into different cortical (i.e., temporal association area) and subcortical areas (i.e., the superior colliculus, periaqueductal gray, posterior thalamic nucleus) simultaneously reveal multiple cell types, namely, intratelencephalic (IT), pyramidal tract (PT) and cortico-thalamic projecting (CT) neurons (Fig. 9C). This connectivity map and derived catalog of anatomically defined neuron types provide complementary and confirmatory information for molecularly defined neuron types described in other BICCN resources (Callaway et al., 2021; Zhang et al., 201; Yao et al., 2021). Finally, these multi-fluorescent retrograde tracing data are available through iConnectome (www.MouseConnectome.org; https://brain.neurobio.ucla.edu/maps/).

**Figure 9.**
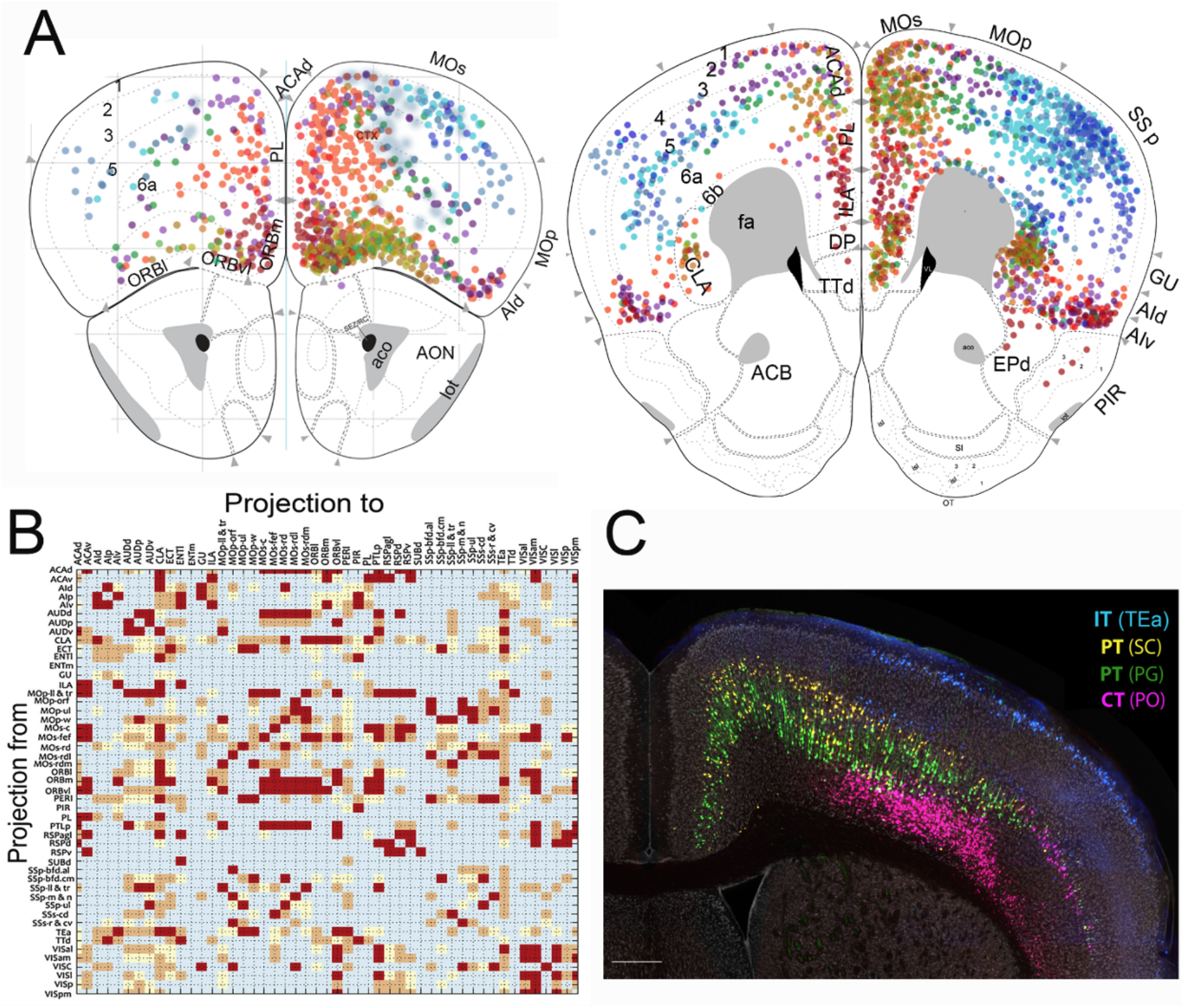
Neuronal Connectivity and Mouse Connectome Project (MCP). A) Connectivity map of distinct cortical projection neurons (cell types) in the prefrontal cortex. **B)** A connectivity matrix constructed based on these retrograde tracing data. These data resources are available on iConnectome www.MouseConnectome.org. C) Example of back labeled neurons in the cortex following injections of four retrograde tracers into the temporal association area (TEa), superior colliculus (SC), periaqueductal gray (PG) and posterior thalamic nucleus (PO), revealing three major classes of cortical neuron types, IT, PT and CT. Scale bar=250μm.

### VI.4 Brain Architecture Project (BAP, RRID:SCR_004283, http://brainarchitecture.org)

High-resolution 2D images from BICCN collaborators are available for display on the Brain Architecture web portal. Datasets are served into species and experiment-type specific pages, accessible from the front landing page (**Suppl. Fig. 4**). Users can filter mouse cell distribution datasets via free text search of metadata for keywords and mouse projection and connectivity datasets via injection region or tracer. A high-resolution viewer capability to display overlays of regional compartments, points indicating cell bodies post cell detection, and skeletonization on 2D sections of atlas mapped brains. The viewer can display data at multiple resolutions, with zoom to super resolution capability, beyond the native in-plane 0.46-μm resolution of the images. All software employed for image analytics pipeline, including registration and atlas mapping, cell detection, and process detection and skeletonization via are available both in interactive versions on the Brain Architecture web portal, and for download (of both source code and documentation) on Github and Bitbucket repositories. There are additionally cross-linkages with the Broad Institute Single-Cell Portal (https://singlecell.broadinstitute.org/single_cell).

### VI.3 Additional Tools and Resources

Numerous other resources have been key in analysis of BICCN publication datasets, described in **Suppl. Materials - BICCN Tools and Resources, Suppl. Table 4**, and at https://biccn.org/tools. Tools key to BICCN publications are *Epiviz* (RRID:SCR_022796), and Brain Cell Methylation Viewer (RRID:SCR_020954), interactive visualization tools for functional genomics data; *Brainome* (RRID:SCR_018162) a genome browser to visualize the cell type-specific transcriptomes and epigenomes of cell types from the mouse MOp; *Catlas* (RRID:SCR_018690), which provides maps of accessible chromatin in the adult mouse isocortex, olfactory bulb, hippocampus and cerebral nuclei; *Cytosplore Viewer* (RRID:SCR_018330) is a stand-alone application (Windows and MacOS) for interactive visual exploration of multi-species and cross-omics single cell data in several BICCN data resources., and MetaNeighbor (Crow et al. 2018; Fischer et al. 2021) a method for assessing the replicability of single cell data, used in a number of key BICCN publications (e.g., (Bakken et al. 2021; Yao, Liu, et al. 2021)) to validate cell types and perform quality control. An important resource for the analysis of single cell brain data is the Broad Institute Single Cell Portal (RRID:SCR_014816).

Several open-access neuroinformatics resources were launched prior to BICCN efforts (Anderson et al. 2021) but were substantially expanded with support and contributions from BICCN projects, and have been utilized in multiple BICCN publications. Two such examples, NeuroMorpho.Org (RRID:SCR_002145) and Hippocampome.org (RRID:SCR_009023), have helped bridge seminal literature information and data with new BICCN-generated data. NeuroMorpho.Org (Akram et al. 2018) provides free access to hundreds of thousands of reconstructed neural cell morphologies contributed by over 900 laboratories worldwide from approximately 100 distinct species, are were utilized in the recent comparative analysis of neocortical neurons (Bakken et al. 2021), where BICCN data from human, marmoset, and mouse were augmented with tracings from other mammals. Hippocampome.org (Hamilton et al. 2017) is a knowledge base of neuron types from the mammalian hippocampal formation and entorhinal cortex with more than 500,000 neuronal properties extracted from 46,000 pieces of evidence annotated from scientific articles. For more details on the above and additional resources, we refer readers to the BICCN online resource Tools and Analysis - Brain Cell Data Center (BCDC).

## VII. Standards and the BICCN: Towards FAIR Neuroscience

To be reused and shared efficiently, accessible data need to be described in standard ways, and the development and adoption of standards is thus essential to advancing rigorous science and efficient collaboration (Poline et al. 2022). An increasingly comprehensive and detailed set of technical, quality control and policy standards developed or utilized by the BICCN provides guidance/best-practices for consortia members and others seeking to use BICCN data. The BICCN is committed to implementing practices and technologies to make data and other research products FAIR (Wilkinson et al., 2016). All data that does not involve protected health information is made available under a CC-BY 4.0 attribution license (Wang et al. 2020).

### VII.1 BICCN Standards, Best Practices and Recommendations

have been implemented across BCDC and the BRAIN data archives including metadata and file formats, common processing pipelines, spatial and semantic standards, and identifier systems. BICCN Working Groups focused on harmonizing protocols, data formats and metadata for transcriptomic, physiological, and anatomical data types. The BCDC coordinated the formation of working groups of consortium members which considered what standards and best practices were necessary for new experimental technologies for which standards were not yet available, including developing QC criteria for a given modality. The BCDC was also responsible for developing strategies to harmonize common metadata across the archives, including submissions checklists, collections metadata and basic descriptive information for specimens. The Metadata and Infrastructure Working Group (**Figure 4F**), comprising representatives from the BCDC, the four BRAIN Archives housing BICCN data and BICCN investigators, coordinated the adoption and development of the necessary technical standards to support FAIR data **(Suppl. Fig. 5**). However, beyond basic descriptive metadata such as modality or species, annotations and mappings at a deeper level are still nascent (Shepherd et al. 2019).

Additional standards were adopted over the course of the project as they became available, e.g., the Essential Metadata for 3D Microscopy standard developed with support from the BRAIN Initiative (Ropelewski et al. 2022) was recently implemented by BIL. The independent data generation within the BICCN allowed post-hoc assessment of standards for rigor and reproducibility via meta-analysis. This is particularly true of the mouse expression data which involved replicates across technologies and allowed assessment and integration to assess the replicability of cell type calling via, e.g. MetaNeighbor (Crow et al. 2018) and post-hoc integration to produce more reliable marker sets (Fischer and Gillis 2021; Bakken et al. 2021). BICCN-developed standards are available through a public GitHub Repository https://github.com/BICCN, and BICCN Standards, Best Practices, and Recommendations - Brain Cell Data Center (BCDC).

### VII.2 BICCN FAIR Data Practices

The BCDC, in partnership with the archives that house the data, ensures that all BICCN data is FAIR (Findable, Accessible, Interoperable and Reusable) according to the principles laid out in (Wilkinson et al. 2016b). The BICCN ecosystem benefits and derives increased utility from the set of 15 FAIR data principles and recommendations. Although full implementation of FAIR was challenging, particularly in the initial phase of the BICCN where the archives, techniques and standards were under simultaneous development, the BICCN has been moving towards implementation of a consistent set of baseline FAIR practices over the course of the project including progress as shown in **Box 1**. This standards-based work includes use of persistent identifiers and rich metadata, detailed provenance, use of FAIR vocabulary, and use of clear data use agreements.

### VII.3 The Brain Data Standards Ontology

An important component of cell type classification is a rigorous and precise ontology and nomenclature. The Brain Data Standards Ontology (BDSO) (Tan et al., n.d.), is an ontology of cell types defined in the BICCN MOp that extends the Cell Ontology (CL) (Diehl et al. 2016) to provide a more detailed set of terms for FAIR-compliant annotation than previously available (Bakken et al., 2017). As an extension of CL, BDSO is fully interoperable with both CL and Uberon (Mungall et al. 2012), allowing data annotated with BDSO terms to be interoperable not only with the BICCN data, but also with datasets from the wider community. This approach is scalable and lowers human error (as compared to manually creating the ontology), features that are crucial in scaling to whole brain annotation. As part of creating BDSO, representation of neuronal cell types in CL has been deepened, adding new cortical cell types by defined markers, projection pattern (e.g. extra telencephalic projecting), layer, and morphology (e.g., pyramidal). These additions to CL have already been used for annotation in datasets in CellXGene (“Chan Zuckerberg CELLxGENE Discover” n.d.), the Cell Annotation Platform (RRID:SCR_022797), and other single cell transcriptomics data providers to deepen annotations to use terms from BDSO. A major application of BDSO is to support organization, navigation and searching of data in the CTKE (**Section VI.1**). Knowledge graphs and APIs were developed for the CTKE, (Knowledge graph: http://purl.obolibrary.org/obo/pcl/bds/kg/; API: http://purl.obolibrary.org/obo/pcl/bds/api/), making the reuse, search, and navigation of the BDSO ontology openly accessible. The latest release of the ontology is hosted at Brain Data Standards Ontology. (**Suppl. Materials - Brain Data Standards Ontology**).

**Box 1.**
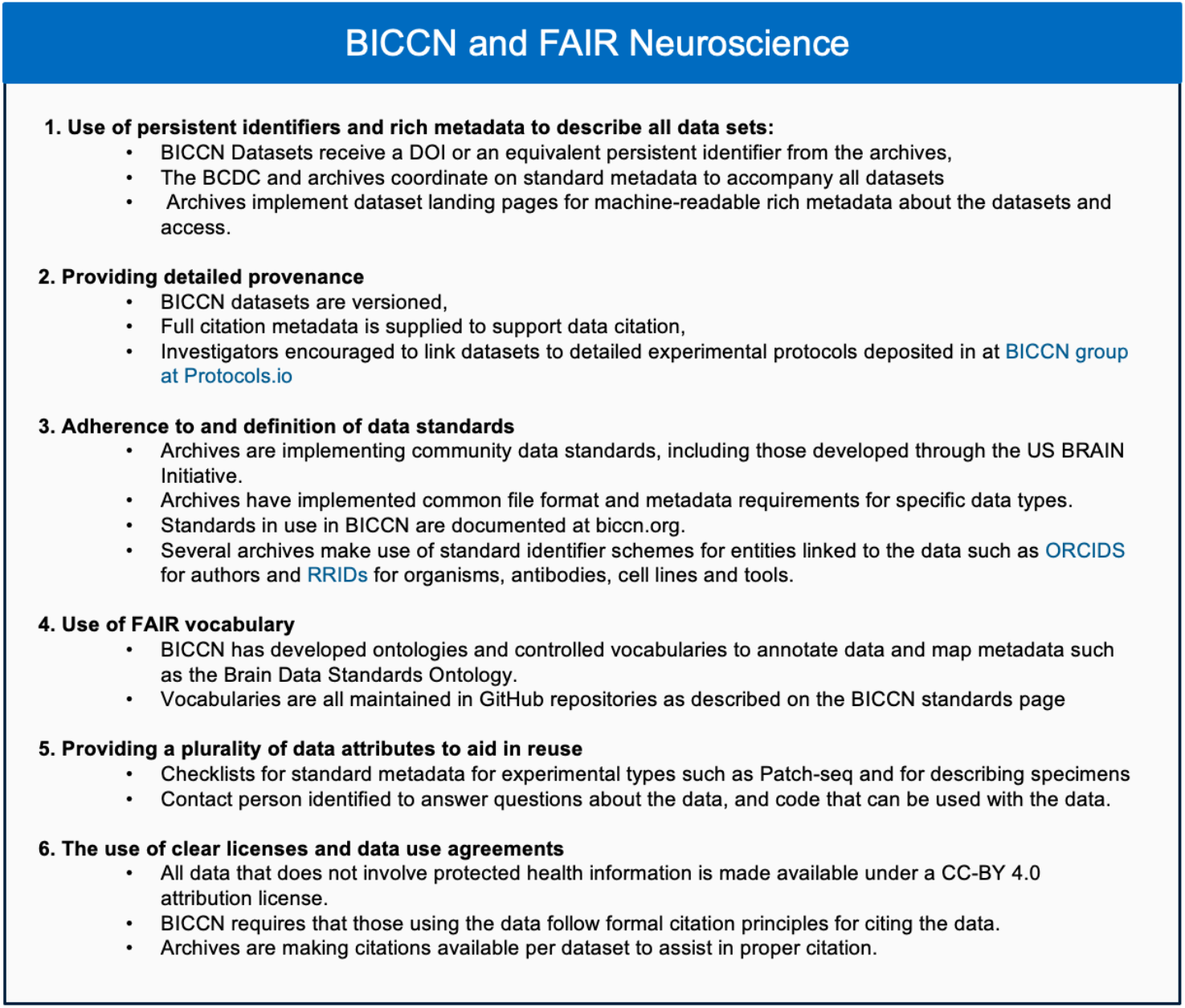
FAIR Neuroscience Data Practices and the BICCN. BICCN data is FAIR (Findable, Accessible, Interoperable and Reusable) according to the principles laid out in (Wilkinson et al. 2016b). The BICCN ecosystem benefits and derives increased utility from the FAIR data principles and recommendations. Summarized are the main areas where the BICCN data ecosystem has implemented these practices.

## VIII. From BICCN to the BRAIN Initiative Cell Atlas Network (BICAN)

Significant progress has been made in the development of laboratory techniques and analysis methods for description of cell types in the mammalian brain. Beginning with the original ten pilot studies of the BICCC in developing, validating, and scaling up emerging genomic and anatomical mapping technologies, the BICCN has used these approaches towards generation of complete, accurate, and permanent (CAP) data resources to form an extensive data ecosystem. At present the BICCN has completed cell type profiling using transcriptomics (10x RNA-seq) and epigenomics (ATAC-seq) for the whole mouse brain, and in many regions of the human brain, and is developing architecture, infrastructure, and product resources to support these data. In addition to ongoing BICCN datasets produced by individual laboratories and resulting publications, **six active BICCN Working Groups** are presently engaged in collaborative projects (BICCN 2.0) integrating and interpreting new and existing data. In addition to fulfilling the goal of integrating transcriptomic and epigenomic data across the entire mouse central nervous system, these groups are developing methods to identify cell-type specific enhancers that can drive systemic delivery of reporter genes to select subclasses or types of brain cells in mice and primates, producing a full molecularly annotated wiring diagrams of the mammalian brain, and beginning preliminary work on developing comprehensive human and NHP atlases, preparing for recently launched Brain Initiative Cell Atlas Network (BICAN). Additional work is in measuring proteomic signatures of brain cells and further developing integration methods and infrastructure for future atlases.

While significant progress has been made in most aspects of the original BICCN infrastructure vision much work remains. The resources developed through the BICCN have been the result of strong collaborations between data generators, analysts, informaticians, and software developers. The ultimately desired data ecosystem will support data collection, quantification, and a mapping framework for managing data and information across diverse repositories. This should be coupled with consistent data description standards that describe and facilitate best practices for community use of multi-modal single cell data and its content. From early in the consortium’s activities, requirements for FAIR data management were discussed, however the goal of building a foundational community resource for housing single-cell centered data content in the brain is still a work in progress.

An important component of full data integration is the spatial mapping of data enabling users to search by spatial location for data of interest, and common coordinate frameworks for mapping must be in place.

There has been general community acceptance of the Allen common coordinate framework for the mouse brain (CCFv3) with several tools now available for pinning specimen level or registering and aligning spatial sections or volumes. While components are now in place for a fully searchable and spatially resolved database, there remains engineering work to incorporate these into a functional application. There also remain challenges in the BICCN infrastructure. The BCDC data catalog offers an entry point to each project and dataset, yet specimen level search and access enumerating regional or nuclei level search is not universally available at present. Further, while the data archives are all capable of accepting and managing large data volumes, and many tools are available for accessing relevant data, this workflow is not yet fully interoperable and there are still inconsistent metadata standards across modalities. This can become challenging for users of multimodality data types such as Patch-seq where the associated data types, transcriptomic, morphology, and electrophysiology are stored in different archives.

BICCN data represent unprecedented coverage describing the cell type landscape of the mammalian brain and the stage is now set for completing the BRAIN Initiative 2025 (“BRAIN 2025 Report” n.d.) vision of large-scale profiling of the human brain including diversity and development. This new phase, commenced in Fall 2022, is the BRAIN Initiative Cell Atlas Network (“RFA-MH-21-235: BRAIN Initiative Cell Atlas Network (BICAN): Comprehensive Center on Human and Non-Human Primate Brain Cell Atlases (UM1 Clinical Trial Not Allowed)” n.d.) and is the extension of the groundwork set by the BICCN. BICAN presents novel challenges of human tissue management and sample selection, the need for improved standardization of sequencing and mapping, and establishment of an integrated neuroinformatics framework. BICAN will present major challenges in establishing standard protocols, mapping, and annotation, but much work can be leveraged from BICCN ecosystem. The neuroinformatics work of the BICAN initiative calls for standardized sequencing and tissue selection, and for the creation of an integrated knowledge base for the community ((“RFA-MH-21-237: BRAIN Initiative Cell Atlas Network (BICAN): Coordinating Unit for Biostatistics, Informatics, and Engagement (CUBIE) (U24 Clinical Trial Not Allowed)” n.d.)).

The ultimate expectation of BRAIN 2025 is to accomplish a full census of neuronal and glial cell types in mouse, human, and non-human primate, an intellectual framework for cell type classification, and to provide experimental access to the different brain cell types to determine their roles in health and disease. However, there is not yet full consensus on what a neuronal type is, since a variety of factors including experience, connectivity, and neuromodulators can diversify the molecular, electrical, and structural properties of initially similar neurons. There is also increasing evidence that there may not even be sharp boundaries separating subtypes from each other, and cell phenotypes may change over time. Here, taxonomies of putative types and representative cells will provide a frame of reference for studies across labs, and possibly in different organisms, allowing cross-comparison.

The extension of the multimodal cell-type atlas of select regions in the human brain to multiple brain regions, particularly those housing vulnerable cell populations, and to different stages of brain development is essential. Such openly available datasets will be key to future studies comparing cell types within their spatial context in the normative brain to those in neuropsychiatric disease, with the addition of transcriptional, epigenetic, morphological, and neurophysiological datasets from postmortem brains, either within the BICCN and BICAN data archives or published in the literature. Combining the BICCN and BICAN data archives with the ability to place cells within a CCF detected from deidentified digital pathology data will also make available large datasets that provide the sample numbers and diverse representation necessary for use of interpretative machine learning analysis applications. Moreover, as techniques for spatial detection of proteins and metabolites achieve multiplexing capabilities as well as cellular resolution, such data may help to uncover disease mechanisms that may be beyond transcriptional and epigenetic detection but, when combined with data currently included in the BICCN datasets, could help explain the neurophysiological changes detected in specific cell types and brain areas as part of a disease phenotype.

The BICCN has provided the community with massive high-quality datasets describing the multimodal cell type landscape of the mammalian brain. Substantial resources now exist for the study of brain cell types, and while the supporting data ecosystem is not yet complete, tremendous progress has been made. Increasingly diverse skills are being applied to the architectural design and development of the new BICAN data ecosystem, and we are planning for continuous extension and enhancement of this work to address human specific challenges. We are only beginning to interpret this valuable data and to understand its importance for the nature of cell types in the brain.

## Supporting information

BICCN Data Ecosystem - Supplementary

Supplementary Table 1_BICCN cell types, modalities, techniques, and investigators

Supplementary Table_2_BICCN Data levels

Supplementary Table_3_ BICCN datasets by levels

Supplementary Table 4_BICCC Data Ecosystem Resources

## BICCN Data Ecosystem Collaboration

**Writing Group.** Michael Hawrylycz^1,*^, Maryann E. Martone^2,3,^*, Patrick R. Hof^4^, Ed Lein^1^, Aviv Regev^5^, Giorgio A. Ascoli^6^, Jan G. Bjaalie^7^, Hong-Wei Dong^8^, Satrajit S. Ghosh^9^, Jesse Gillis^10^, Ronna Hertzano^11,12,13^, David R Haynor^14^, Yongsoo Kim^15^, Yufeng Liu^16^, Jeremy A Miller^1^, Partha P. Mitra^17^, Eran Mukamel^18^, David Osumi-Sutherland^19^, Hanchuan Peng^16^, Patrick L. Ray^1^, Raymond Sanchez^1^, Alex Ropelewski^28^, Richard H. Scheuermann^20^, Shawn Zheng Kai Tan^19^, Timothy Tickle^21^, Hagen Tilgner^22^, Merina Varghese^4^, Brock Wester^23^, Owen White^13^

**BICCN Ecosystem Scientific Oversight.** Michael Hawrylycz^1^, Lydia Ng^1^, Carol L. Thompson^1^, Maryann E. Martone^2,3^,, Giorgio A. Ascoli^6^, Joseph R. Ecker^24,25^, Satrajit Ghosh^9^, Ed Lein^1^, Eran Mukamel^18^, Bing Ren^26,27^, Alex Roplewski^28^, Owen White^13^, Hongkui Zeng^1^

**BICCN Infrastructure Working Group.** Carol L. Thompson^1^, Michael Hawrylycz^1^, Pamela M. Baker^1^, Anita Bandrowski^2^, Heather Creasy^13^, Satrajit S. Ghosh^9^, Tom Gillespie^2^, Gregory Hood^28^, Maryann E. Martone^2,3^, Anup Markuhar^13^, Lydia Ng^1^, Timothy Tickle^21^, Alex Ropelewski^28^, Owen White^13^, Patrick L. Ray^1^, Hua Xu^29^, W. Jim Zheng^29^, Guo-Qiang Zhang^30^

**BICCN Portal.** Carol L. Thompson^1^, Michael Hawrylycz^1^, Maryann E. Martone^2,3^, Anita Bandrowski^2^, Tim Dolbeare^1^, Shoaib Mufti^1^, Lydia Ng^1^, Patrick L. Ray^1^, Brian Staats^1^

**BICCN Tools and Resources.** Brian Aevermann^31^, Cody Baker^37^, Anita Bandrowski^2^, Ambrose Carr^31^, Roni Choudhury^38^, Jonah Cool^31^, Tim Dolbeare^1^, Jesse Gillis^10^, Nathan Gouwens^1^, Boudewijn Lelieveldt^32,33^, Hanqing Liu^24^, Eran A. Mukamel^18^, David Osumi-Sutherland^19^, Hanchuan Peng^16^, Raymond Sanchez^1^, Richard H. Scheuermann^20^, Tim Tickle^21^, Hagen Tilgner^22^, Rongxin Fang^34,35^, Yang Li^26^, Fangming Xie^36^, Yun (Renee) Zhang^20^,

**BICCN Data Levels Group.** Michael Hawrylycz^1^, Maryann E. Martone^2,3^, Carol L. Thompson^1^

**Common Coordinate Frameworks and Image Registration.** Lydia Ng^1^, James Gee^41^,, David Allemang^38^, Thomas L. Athey^40^, Jan G. Bjaalie^7^, Min Chen^41^, Roni Choudhury^38^, Song-Lin Ding^1^, Jean-Christophe Fillion-Robin^38^, Nathan Gouwens^1^, Sam Horvath^38^, Shengdian Jiang^16^, Yufeng Liu^16^, Lijuan Liu^16^, Michael I. Miller^40^, Hanchuan Peng^16^,, Maja A Puchades^7^, Daniel Tward^40^, Lei Qu^16^, Quanxin Wang^1^, Zhixi Yun^16^

**BRAIN Cell Data Center (BCDC) Consortium administration and Data publishing.** Carol L. Thompson^1^, Pamela M. Baker^1^, Prajal Bishwakarma^1^, Timothy P. Fliss^1^, Katherine S. Baker^1^, Florence D’Orazi^31^, Lauren Kruse^1^, James Mathews^1^, Nathan Sjoquist^42^, Lydia Ng^1^, Michael Hawrylycz^1^

**Molecular Pipelines Group.** Timothy Tickle^21^, Zizhen Yao^1^, Heather Creasy^13^, Kylee Degatano^21^, Brian R. Herb^13^, Elizabeth A. Kiernan^21^, Farzaneh Khajouei^21^, Changkyu Lee^1^, Hanqing Liu^24^, Kaylee L. Mathews^21^, Eran Mukamel^18^, Hagen Tilgner^22^, Cindy T.J. van Velthoven^1^, Fangming Xie^36^, Owen White^13^

**NeMO Archive.** Owen White^13^, Heather Creasy^13^, Anup Mahurkar^13^

**DANDI Archive.** Satrajit S. Ghosh^9^, Cody Baker^37^, Roni Choudhury^38^, Benjamin Dichter^37^, Yaroslav O. Halchenko^39^, Dorota Jarecka^9^

**BIL Archive.** Alexander J Ropelewski^28^, Gregory Hood^28^

**BossDB Archive.** Brock Wester^23^

**SEU-Allen Center.** Hanchuan Peng^16^,, Lijuan Liu^16^,, Yufeng Liu^16^, Lei Qu^16^, Zhixi Yun^16^, Shengdian Jiang^16^

**Brain Architecture Portal.** Partha P. Mitra^17^, Samik Banerjee^17^, Bingxing Huo^17^, Chris Mezias^17^,, Daniel Tward^40^

**Cell Type Knowledge Explorer.** Raymond Sanchez^1^, Jeremy A. Miller^1^, Patrick L. Ray,^1^ Tim Dolbeare^1^, Brian Aevermann^31^, Michael Hawrylycz^1^, Huseyin Kir^19^, Ed Lein^1^, Tyler Mollenkopf^1^, David Osumi-Sutherland^19^, Shoaib Mufti^1^, Richard H. Scheuermann^20^, Brian Staats^1^, Shawn Zheng Kai Tan^19^, Hongkui Zeng^1^, Yun (Renee) Zhang^20^

**NeMO Analytics.** Ronna Hertzano^11,12,13^, Seth Ament^13^, Brian R. Herb^13^, Joshua Orvis^13^, Joseph P. Receveur^13^

**Mouse Connectome Project.** Hong-Wei Dong^8^, Houri Hintiryan^8^, Brian Zingg^8^, Giorgio Ascoli^6^

**Brain Data Standards Ontology.** David Osumi-Sutherland^19^, Patrick L. Ray^1^, Brian Aevermann^31^, Tom Gillespie^2^, Nomi L. Harris^43^, Huseyin Kir^19^, Ed Lein^1^, Jeremy A. Miller^1^, Christopher J. Mungall^43^, Raymond Sanchez^1^, Shawn Zheng Kai Tan^19^, Richard H. Scheuermann^20^, Yun (Renee) Zhang^20^

**FAIR Data Standards.** Maryann E. Martone^2,3^, Anita Bandrowski^2^, Michael Hawrylycz^1^, Carol L. Thompson^1^

^1^Allen Institute for Brain Science, Seattle, WA

^2^Dept of Neuroscience, University of California San Diego, San Diego, CA

^3^San Francisco VA Medical Center, San Francisco, CA

^4^Nash Family Department of Neuroscience and Friedman Brain Institute, Icahn School of Medicine at Mount Sinai, New York, NY

^5^Broad Institute of MIT and Harvard, Cambridge, Current address: Genentech, 1 DNA Way, South, San Francisco, CA

^6^Bioengineering Department and Center for Neural Informatics, Structures, & Plasticity, Volgenau School of Engineering, George Mason University, Fairfax, VA

^7^Institute of Basic Medical Sciences, University of Oslo, Oslo, Norway

^8^ UCLA Brain Research & Artificial Intelligence Nexus, Department of Neurobiology, David Geffen School of Medicine at UCLA. 73-235 CHS, Los Angeles, CA

^9^McGovern Institute for Brain Research, Massachusetts Institute of Technology, Cambridge, MA

^10^Department of Physiology, University of Toronto, Toronto, Canada

^11^Department of Otorhinolaryngology Head and Neck Surgery, University of Maryland School of Medicine, Baltimore, MD

^12^Department of Anatomy and Neurobiology, University of Maryland School of Medicine, Baltimore MD

^13^Institute for Genome Sciences, University of Maryland School of Medicine, Baltimore, MD

^14^Department of Radiology, University of Washington, Seattle, WA

^15^Department of Neural and Behavioral Sciences, College of Medicine, The Pennsylvania State University, PA

^16^SEU-Allen Institute Joint Center, Institute for Brain and Intelligence, Southeast University, Nanjing, China

^17^Cold Spring Harbor Laboratory, Cold Spring Harbor, New York, NY

^18^Department of Cognitive Science, University of California, San Diego, La Jolla, CA

^19^European Bioinformatics Institute (EMBL-EBI), Wellcome Trust Genome Campus, Hinxton, Cambridge, United Kingdom

^20^J. Craig Venter Institute, La Jolla, CA, USA, Department of Pathology, University of California San Diego, La Jolla, CA

^21^Data Sciences Platform, Broad Institute of MIT and Harvard, Cambridge, MA

^22^Feil Family Brain and Mind Research Institute, Weill Cornell Medicine, NY, USA. Center for Neurogenetics, Weill Cornell Medicine, NY

^23^Research and Exploratory Development Department, Johns Hopkins University Applied Physics Laboratory, Laurel, MD

^24^Genomic Analysis Laboratory, The Salk Institute for Biological Studies, La Jolla, CA

^25^Howard Hughes Medical Institute, The Salk Institute for Biological Studies, La Jolla, CA

^26^Center for Epigenomics, Department of Cellular and Molecular Medicine, UC San Diego School of Medicine, La Jolla, CA

^27^Ludwig Institute for Cancer Research, La Jolla, CA

^28^Pittsburgh Supercomputing Center, Carnegie Mellon University, Pittsburgh, PA

^29^School of Biomedical Informatics, The University of Texas Health Science Center at Houston, Houston, TX

^30^ Texas Institute for Restorative Neurotechnologies. The University of Texas Health Science Center at Houston, Houston, TX

^31^Chan Zuckerberg Initiative, Redwood City, CA

^32^Department. of Radiology, Leiden University Medical Center, Leiden, NL,

^33^Dept. of Intelligent Systems, Faculty of EEMCS, Delft University of Technology, Delft, NL

^34^Ludwig Institute for Cancer Research, La Jolla, CA,

^35^Bioinformatics and Systems Biology Graduate Program, University of California San Diego, La Jolla, CA

^36^Department of Chemistry and Biochemistry, University of California Los Angeles, CA

^37^CatalystNeuro, Benicia, CA

^38^Kitware Inc., Albany, NY

^39^Department of Psychological and Brain Sciences, Dartmouth College, Hannover, NH

^40^Department of Biomedical Engineering, Johns Hopkins University, Baltimore, MD

^41^Department of Radiology, Perelman School of Medicine, University of Pennsylvania, Philadelphia, PA

^42^Microsoft, Seattle, WA

^43^Environmental Genomics and Systems Biology Division, Lawrence Berkeley National Laboratory, Berkeley, CA

*Correspondence

## Acknowledgments

Research reported in this publication was supported by the National Institute of Mental Health of the National Institutes of Health. The content is solely the responsibility of the authors and does not necessarily represent the official views of the National Institutes of Health. MJH, MEM, CLT, JG were supported by NIH grants U19MH114824, U19MH114824, R01MH123220. The Allen Institute founder, Paul G. Allen, for his vision and support. GAA was supported in part by NIH grants R01NS39600, U01MH114829, RF1MH128693, and R01NS86082, and by DOE grant DE-SC0022998. BW thanks Erik C. Johnson’s work in updating the metadata standard/service for BossDB. JGB was supported for the QuickNII and VisuAlign tools by the European Union’s Horizon 2020 Framework Programme for Research and Innovation under the Framework Partnership Agreement No. 650003 (HBP FPA). YL and LL acknowledge funding from Southeast University (SEU) to support informatics data management and analysis pipeline of full neuronal reconstruction platform, and MOST (China) Brain Research Project, “Mammalian Whole Brain Mesoscopic Stereotaxic 3D Atlas” (2022ZD0205200, 2022ZD0205204). PRH was supported byU01 MH117023, JG R01 NS096720, Yongsoo Kim by NIH RF1MH124605 Brain Data Standards Ontology team was supported in part by NIH grants 1R01MH123220-01 and 1RF1MH123220-01, CJM and NLH were supported in part by the Director, Office of Science, Office of Basic Energy Sciences of the U.S. Department of Energy Contract No. DE-AC02-05CH11231. BW acknowledges R24MH114785 and thanks to Erik C. Johnson. The DANDI team acknowledges support from R24MH117295. The BICCN Data Ecosystem Collaboration is grateful for the vision and support of NIH BRAIN Initiative Director John Ngai, and NIMH program staff Yong Yao, Ming Zhan, and Laura Reyes.

## References

Aevermann, Brian, Yun Zhang, Mark Novotny, Mohamed Keshk, Trygve Bakken, Jeremy Miller, Rebecca Hodge, Boudewijn Lelieveldt, Ed Lein, and Richard H. Scheuermann. 2021. “A Machine Learning Method for the Discovery of Minimum Marker Gene Combinations for Cell Type Identification from Single-Cell RNA Sequencing.” Genome Research 31 (10): 1767–80.

Akram, Masood A., Sumit Nanda, Patricia Maraver, Rubén Armañanzas, and Giorgio A. Ascoli. 2018. “An Open Repository for Single-Cell Reconstructions of the Brain Forest.” Scientific Data 5 (February): 180006.

Anderson, Kristin R., Julie A. Harris, Lydia Ng, Pjotr Prins, Sara Memar, Bengt Ljungquist, Daniel Fürth, Robert W. Williams, Giorgio A. Ascoli, and Dani Dumitriu. 2021. “Highlights from the Era of Open Source Web-Based Tools.” The Journal of Neuroscience: The Official Journal of the Society for Neuroscience 41 (5): 927–36.

Avants, B. B., C. L. Epstein, M. Grossman, and J. C. Gee. 2008. “Symmetric Diffeomorphic Image Registration with Cross-Correlation: Evaluating Automated Labeling of Elderly and Neurodegenerative Brain.” Medical Image Analysis 12 (1): 26–41.

Bakken, Trygve E., Nikolas L. Jorstad, Qiwen Hu, Blue B. Lake, Wei Tian, Brian E. Kalmbach, Megan Crow, et al. 2021. “Comparative Cellular Analysis of Motor Cortex in Human, Marmoset and Mouse.” Nature 598 (7879): 111–19.

Baskarada, Sasa, and Andy Koronios. 2013. “Australasian Journal of Information Systems.” Australasian Journal of Information Systems 18 (1). https://doi.org/10.3127/ajis.v18i1.748.

Benavidez, Nora L., Michael S. Bienkowski, Muye Zhu, Luis H. Garcia, Marina Fayzullina, Lei Gao, Ian Bowman, et al. 2021a. “Organization of the Inputs and Outputs of the Mouse Superior Colliculus.” Nature Communications 12 (1): 4004.

Benavidez, Nora L., Michael S. Bienkowski, Muye Zhu, Luis H. Garcia, Marina Fayzullina, Lei Gao, Ian Bowman, et al. 2021b. “Organization of the Inputs and Outputs of the Mouse Superior Colliculus.” Nature Communications 12 (1): 4004.

“BICCN Omics Workshop 2022.” n.d. Accessed September 20, 2022a. https://nemoarchive.org/biccn-omics-workshop/.

“BICCN Omics Workshop 2022.” n.d. Accessed September 20, 2022b. https://nemoarchive.org/biccn-omics-workshop/.

Bienkowski, Michael S., Ian Bowman, Monica Y. Song, Lin Gou, Tyler Ard, Kaelan Cotter, Muye Zhu, et al. 2018. “Integration of Gene Expression and Brain-Wide Connectivity Reveals the Multiscale Organization of Mouse Hippocampal Networks.” Nature Neuroscience 21 (11): 1628–43.

Booeshaghi, A. Sina, Zizhen Yao, Cindy van Velthoven, Kimberly Smith, Bosiljka Tasic, Hongkui Zeng, and Lior Pachter. 2021. “Isoform Cell-Type Specificity in the Mouse Primary Motor Cortex.” Nature 598 (7879): 195–99.

“BRAIN 2025 Report.” n.d. Accessed March 31, 2022. https://braininitiative.nih.gov/strategic-planning/brain-2025-report.

“Brain Image Library: Contact.” n.d. Accessed September 20, 2022a. https://www.brainimagelibrary.org/training.html.

“Brain Image Library: Contact.” n.d. Accessed September 20, 2022b. https://www.brainimagelibrary.org/training.html.

BRAIN Initiative Cell Census Network (BICCN). 2021. “A Multimodal Cell Census and Atlas of the Mammalian Primary Motor Cortex.” Nature 598 (7879): 86–102.

Chandrashekhar, Vikram, Daniel J. Tward, Devin Crowley, Ailey K. Crow, Matthew A. Wright, Brian Y. Hsueh, Felicity Gore, et al. 2021. “CloudReg: Automatic Terabyte-Scale Cross-Modal Brain Volume Registration.” Nature Methods 18 (8): 845–46.

“Chan Zuckerberg CELLxGENE Discover.” n.d. Cellxgene Data Portal. Accessed September 22, 2022. https://cellxgene.cziscience.com/.

Crow, Megan, Anirban Paul, Sara Ballouz, Z. Josh Huang, and Jesse Gillis. 2018. “Characterizing the Replicability of Cell Types Defined by Single Cell RNA-Sequencing Data Using MetaNeighbor.” Nature Communications 9 (1): 1–12.

Di Bella, Daniela J., Ehsan Habibi, Robert R. Stickels, Gabriele Scalia, Juliana Brown, Payman Yadollahpour, Sung Min Yang, et al. 2021. “Molecular Logic of Cellular Diversification in the Mouse Cerebral Cortex.” Nature 595 (7868): 554–59.

Diehl, Alexander D., Terrence F. Meehan, Yvonne M. Bradford, Matthew H. Brush, Wasila M. Dahdul, David S. Dougall, Yongqun He, et al. 2016. “The Cell Ontology 2016: Enhanced Content, Modularization, and Ontology Interoperability.” Journal of Biomedical Semantics 7 (1): 44.

Ding, Song-Lin, Joshua J. Royall, Susan M. Sunkin, Lydia Ng, Benjamin A. C. Facer, Phil Lesnar, Angie Guillozet-Bongaarts, et al. 2017. “Comprehensive Cellular-Resolution Atlas of the Adult Human Brain.” The Journal of Comparative Neurology 525 (2): 407.

Ecker, Joseph R., Daniel H. Geschwind, Arnold R. Kriegstein, John Ngai, Pavel Osten, Damon Polioudakis, Aviv Regev, Nenad Sestan, Ian R. Wickersham, and Hongkui Zeng. 2017. “The BRAIN Initiative Cell Census Consortium: Lessons Learned toward Generating a Comprehensive Brain Cell Atlas.” Neuron 96 (3): 542–57.

Fedorov, Andriy, Reinhard Beichel, Jayashree Kalpathy-Cramer, Julien Finet, Jean-Christophe Fillion-Robin, Sonia Pujol, Christian Bauer, et al. 2012. “3D Slicer as an Image Computing Platform for the Quantitative Imaging Network.” Magnetic Resonance Imaging 30 (9): 1323–41.

Fischer, Stephan, Megan Crow, Benjamin D. Harris, and Jesse Gillis. 2021. “Scaling up Reproducible Research for Single-Cell Transcriptomics Using MetaNeighbor.” Nature Protocols 16 (8): 4031–67.

Fischer, Stephan, and Jesse Gillis. 2021. “How Many Markers Are Needed to Robustly Determine a Cell’s Type?” iScience 24 (11): 103292.

Foster, Nicholas N., Joshua Barry, Laura Korobkova, Luis Garcia, Lei Gao, Marlene Becerra, Yasmine Sherafat, et al. 2021. “The Mouse Cortico-Basal Ganglia-Thalamic Network.” Nature 598 (7879): 188–94.

Gao, Le, Sang Liu, Lingfeng Gou, Yachuang Hu, Yanhe Liu, Li Deng, Danyi Ma, et al. 2022. “Single-Neuron Projectome of Mouse Prefrontal Cortex.” Nature Neuroscience 25 (4): 515–29.

Gorgolewski, Krzysztof J., Tibor Auer, Vince D. Calhoun, R. Cameron Craddock, Samir Das, Eugene P. Duff, Guillaume Flandin, et al. 2016. “The Brain Imaging Data Structure, a Format for Organizing and Describing Outputs of Neuroimaging Experiments.” Scientific Data 3 (June): 160044.

Halchenko, Yaroslav, Kyle Meyer, Benjamin Poldrack, Debanjum Solanky, Adina Wagner, Jason Gors, Dave MacFarlane, et al. 2021. “DataLad: Distributed System for Joint Management of Code, Data, and Their Relationship.” Journal of Open Source Software. https://doi.org/10.21105/joss.03262.

Hamilton, D. J., D. W. Wheeler, C. M. White, C. L. Rees, A. O. Komendantov, M. Bergamino, and G. A. Ascoli. 2017. “Name-Calling in the Hippocampus (and Beyond): Coming to Terms with Neuron Types and Properties.” Brain Informatics 4 (1): 1–12.

Hao, Yuhan, Stephanie Hao, Erica Andersen-Nissen, William M. Mauck 3rd, Shiwei Zheng, Andrew Butler, Maddie J. Lee, et al. 2021. “Integrated Analysis of Multimodal Single-Cell Data.” Cell 184 (13): 3573–87.e29.

Hardwick, Simon A., Wen Hu, Anoushka Joglekar, Li Fan, Paul G. Collier, Careen Foord, Jennifer Balacco, et al. 2022. “Single-Nuclei Isoform RNA Sequencing Unlocks Barcoded Exon Connectivity in Frozen Brain Tissue.” Nature Biotechnology 40 (7): 1082–92.

Hawrylycz, Michael J., Ed S. Lein, Angela L. Guillozet-Bongaarts, Elaine H. Shen, Lydia Ng, Jeremy A. Miller, Louie N. van de Lagemaat, et al. 2012. “An Anatomically Comprehensive Atlas of the Adult Human Brain Transcriptome.” Nature 489 (7416): 391–99.

Hodge, Rebecca D., Trygve E. Bakken, Jeremy A. Miller, Kimberly A. Smith, Eliza R. Barkan, Lucas T. Graybuck, Jennie L. Close, et al. 2019. “Conserved Cell Types with Divergent Features in Human versus Mouse Cortex.” Nature 573 (7772): 61–68.

HuBMAP Consortium. 2019. “The Human Body at Cellular Resolution: The NIH Human Biomolecular Atlas Program.” Nature 574 (7777): 187–92.

Insel, T. R., S. C. Landis, and F. S. Collins. 2013. “The NIH BRAIN Initiative.” Science. https://doi.org/10.1126/science.1239276.

Jiang, Shengdian, Yimin Wang, Lijuan Liu, Liya Ding, Zongcai Ruan, Hong-Wei Dong, Giorgio A. Ascoli, Michael Hawrylycz, Hongkui Zeng, and Hanchuan Peng. 2022. “Petabyte-Scale Multi-Morphometry of Single Neurons for Whole Brains.” Neuroinformatics, February. https://doi.org/10.1007/s12021-022-09569-4.

Klein, Stefan, Marius Staring, Keelin Murphy, Max A. Viergever, and Josien P. W. Pluim. 2010. “Elastix: A Toolbox for Intensity-Based Medical Image Registration.” IEEE Transactions on Medical Imaging 29 (1): 196–205.

Kozareva, Velina, Caroline Martin, Tomas Osorno, Stephanie Rudolph, Chong Guo, Charles Vanderburg, Naeem Nadaf, Aviv Regev, Wade G. Regehr, and Evan Macosko. 2021. “A Transcriptomic Atlas of Mouse Cerebellar Cortex Comprehensively Defines Cell Types.” Nature 598 (7879): 214–19.

Liu, Hanqing, Jingtian Zhou, Wei Tian, Chongyuan Luo, Anna Bartlett, Andrew Aldridge, Jacinta Lucero, et al. 2021. “DNA Methylation Atlas of the Mouse Brain at Single-Cell Resolution.” Nature 598 (7879): 120–28.

Li, Yang Eric, Sebastian Preissl, Xiaomeng Hou, Ziyang Zhang, Kai Zhang, Yunjiang Qiu, Olivier B. Poirion, et al. 2021. “An Atlas of Gene Regulatory Elements in Adult Mouse Cerebrum.” Nature 598 (7879): 129–36.

Li, Yuanyuan, Jun Wu, Donghuan Lu, Chao Xu, Yefeng Zheng, Hanchuan Peng, and Lei Qu. 2022. “mBrainAligner-Web: A Web Server for Cross-Modal Coherent Registration of Whole Mouse Brains.” Bioinformatics, August. https://doi.org/10.1093/bioinformatics/btac549.

Mai, Juergen K., and Milan Majtanik. 2017. Human Brain in Standard MNI Space: A Comprehensive Pocket Atlas. Elsevier.

Matho, Katherine S., Dhananjay Huilgol, William Galbavy, Miao He, Gukhan Kim, Xu An, Jiangteng Lu, et al. 2021. “Genetic Dissection of the Glutamatergic Neuron System in Cerebral Cortex.” Nature. https://doi.org/10.1038/s41586-021-03955-9.

Merelli, Ivan, Horacio Pérez-Sánchez, Sandra Gesing, and Daniele D’Agostino. 2014. “Managing, Analysing, and Integrating Big Data in Medical Bioinformatics: Open Problems and Future Perspectives.” BioMed Research International 2014 (September): 134023.

Mukamel, Eran A., and John Ngai. 2019. “Perspectives on Defining Cell Types in the Brain.” Current Opinion in Neurobiology 56 (June): 61–68.

Mungall, Christopher J., Carlo Torniai, Georgios V. Gkoutos, Suzanna E. Lewis, and Melissa A. Haendel. 2012. “Uberon, an Integrative Multi-Species Anatomy Ontology.” Genome Biology 13 (1): R5.

Muñoz-Castañeda, Rodrigo, Brian Zingg, Katherine S. Matho, Xiaoyin Chen, Quanxin Wang, Nicholas N. Foster, Anan Li, et al. 2021. “Cellular Anatomy of the Mouse Primary Motor Cortex.” Nature 598 (7879): 159–66.

“NeuroMorpho.Org - a Centrally Curated Inventory of Digitally Reconstructed Neurons and Glia.” n.d. Accessed September 20, 2022. https://www.neuromorpho.org.

“NIH Strategic Plan for Data Science.” n.d. Accessed September 20, 2022. https://datascience.nih.gov/nih-strategic-plan-data-science.

Peng, Hanchuan, Zongcai Ruan, Fuhui Long, Julie H. Simpson, and Eugene W. Myers. 2010. “V3D Enables Real-Time 3D Visualization and Quantitative Analysis of Large-Scale Biological Image Data Sets.” Nature Biotechnology 28 (4): 348–53.

Peng, Hanchuan, Peng Xie, Lijuan Liu, Xiuli Kuang, Yimin Wang, Lei Qu, Hui Gong, et al. 2021. “Morphological Diversity of Single Neurons in Molecularly Defined Cell Types.” Nature 598 (7879): 174–81.

Poline, Jean-Baptiste, David N. Kennedy, Friedrich T. Sommer, Giorgio A. Ascoli, David C. Van Essen, Adam R. Ferguson, Jeffrey S. Grethe, et al. 2022. “Is Neuroscience FAIR? A Call for Collaborative Standardisation of Neuroscience Data.” Neuroinformatics. https://doi.org/10.1007/s12021-021-09557-0.

Puchades, Maja A., Gergely Csucs, Debora Ledergerber, Trygve B. Leergaard, and Jan G. Bjaalie. 2019. “Spatial Registration of Serial Microscopic Brain Images to Three-Dimensional Reference Atlases with the QuickNII Tool.” PloS One 14 (5): e0216796.

Qu, Lei, Yuanyuan Li, Peng Xie, Lijuan Liu, Yimin Wang, Jun Wu, Yu Liu, et al. 2022. “Cross-Modal Coherent Registration of Whole Mouse Brains.” Nature Methods 19 (1): 111–18.

Regev, Aviv, Sarah A. Teichmann, Eric S. Lander, Ido Amit, Christophe Benoist, Ewan Birney, Bernd Bodenmiller, et al. 2017. “The Human Cell Atlas.” eLife 6 (December). https://doi.org/10.7554/eLife.27041.

“RFA-MH-21-235: BRAIN Initiative Cell Atlas Network (BICAN): Comprehensive Center on Human and Non-Human Primate Brain Cell Atlases (UM1 Clinical Trial Not Allowed).” n.d. Accessed August 18, 2022a. https://grants.nih.gov/grants/guide/rfa-files/RFA-MH-21-235.html.

“RFA-MH-21-235: BRAIN Initiative Cell Atlas Network (BICAN): Comprehensive Center on Human and Non-Human Primate Brain Cell Atlases (UM1 Clinical Trial Not Allowed).” n.d. Accessed July 14, 2022b. https://grants.nih.gov/grants/guide/rfa-files/RFA-MH-21-235.html. “RFA-MH-21-237: BRAIN Initiative Cell Atlas Network (BICAN): Coordinating Unit for Biostatistics,

Informatics, and Engagement (CUBIE) (U24 Clinical Trial Not Allowed).” n.d. Accessed September 23, 2022. https://grants.nih.gov/grants/guide/rfa-files/RFA-MH-21-237.html.

Ropelewski, Alexander J., Megan A. Rizzo, Jason R. Swedlow, Jan Huisken, Pavel Osten, Neda Khanjani, Kurt Weiss, et al. 2022. “Standard Metadata for 3D Microscopy.” Scientific Data. https://doi.org/10.1038/s41597-022-01562-5.

Rübel, Oliver, Andrew Tritt, Ryan Ly, Benjamin K. Dichter, Satrajit Ghosh, Lawrence Niu, Pamela Baker, et al. 2022. “The Neurodata Without Borders Ecosystem for Neurophysiological Data Science.” eLife 11 (October). https://doi.org/10.7554/eLife.78362.

Safran, Marilyn, Irina Dalah, Justin Alexander, Naomi Rosen, Tsippi Iny Stein, Michael Shmoish, Noam Nativ, et al. 2010. “GeneCards Version 3: The Human Gene Integrator.” Database: The Journal of Biological Databases and Curation 2010 (August): baq020.

Scala, Federico, Dmitry Kobak, Matteo Bernabucci, Yves Bernaerts, Cathryn René Cadwell, Jesus Ramon Castro, Leonard Hartmanis, et al. 2020. “Phenotypic Variation of Transcriptomic Cell Types in Mouse Motor Cortex.” Nature 598 (7879): 144–50.

Schindelin, Johannes, Ignacio Arganda-Carreras, Erwin Frise, Verena Kaynig, Mark Longair, Tobias Pietzsch, Stephan Preibisch, et al. 2012. “Fiji: An Open-Source Platform for Biological-Image Analysis.” Nature Methods 9 (7): 676–82.

Shepherd, Gordon M., Luis Marenco, Michael L. Hines, Michele Migliore, Robert A. McDougal, Nicholas T. Carnevale, Adam J. H. Newton, Monique Surles-Zeigler, and Giorgio A. Ascoli. 2019. “Neuron Names: A Gene- and Property-Based Name Format, With Special Reference to Cortical Neurons.” Frontiers in Neuroanatomy 13 (March): 25.

Singh, Prachi. 2016. “Big Genomic Data in Bioinformatics Cloud.” Applied Microbiology: Open Access. https://doi.org/10.4172/2471-9315.1000113.

Sontheimer, Harald. 2021. Diseases of the Nervous System. Elsevier.

Tan, Shawn Zheng Kai, Huseyin Kir, Brian D. Aevermann, Tom Gillespie, Nomi Harris, Michael Hawrylycz, Nik Jorstad, et al. n.d. “Brain Data Standards - A Method for Building Data-Driven Cell-Type Ontologies.” https://doi.org/10.1101/2021.10.10.463703.

Wang, Q., S. L. Ding, Y. Li, J. Royall, D. Feng, P. Lesnar, N. Graddis, et al. 2020. “The Allen Mouse Brain Common Coordinate Framework: A 3D Reference Atlas.” Cell 181 (4). https://doi.org/10.1016/j.cell.2020.04.007.

Wilkinson, Mark D., Michel Dumontier, I. Jsbrand Jan Aalbersberg, Gabrielle Appleton, Myles Axton, Arie Baak, Niklas Blomberg, et al. 2016a. “The FAIR Guiding Principles for Scientific Data Management and Stewardship.” Scientific Data 3 (March): 160018.

Wilkinson, Mark D., Michel Dumontier, I. Jsbrand Jan Aalbersberg, Gabrielle Appleton, Myles Axton, Arie Baak, Niklas Blomberg, et al. 2016b. “The FAIR Guiding Principles for Scientific Data Management and Stewardship.” Scientific Data 3 (March): 160018.

Winnubst, Johan, Erhan Bas, Tiago A. Ferreira, Zhuhao Wu, Michael N. Economo, Patrick Edson, Ben J. Arthur, et al. 2019. “Reconstruction of 1,000 Projection Neurons Reveals New Cell Types and Organization of Long-Range Connectivity in the Mouse Brain.” Cell 179 (1): 268–81.e13.

Yao, Zizhen, Hanqing Liu, Fangming Xie, Stephan Fischer, Ricky S. Adkins, Andrew I. Aldridge, Seth A. Ament, et al. 2021. “A Transcriptomic and Epigenomic Cell Atlas of the Mouse Primary Motor Cortex.” Nature 598 (7879): 103–10.

Yao, Zizhen, Cindy T. J. van Velthoven, Thuc Nghi Nguyen, Jeff Goldy, Adriana E. Sedeno-Cortes, Fahimeh Baftizadeh, Darren Bertagnolli, et al. 2021. “A Taxonomy of Transcriptomic Cell Types across the Isocortex and Hippocampal Formation.” Cell 184 (12): 3222–41.e26.

Yuste, Rafael, Michael Hawrylycz, Nadia Aalling, Argel Aguilar-Valles, Detlev Arendt, Ruben Armañanzas, Giorgio A. Ascoli, et al. 2020. “A Community-Based Transcriptomics Classification and Nomenclature of Neocortical Cell Types.” Nature Neuroscience 23 (12): 1456–68.

Zeng, Hongkui. 2022. “What Is a Cell Type and How to Define It?” Cell 185 (15): 2739–55.

Zhang, Meng, Stephen W. Eichhorn, Brian Zingg, Zizhen Yao, Kaelan Cotter, Hongkui Zeng, Hongwei Dong, and Xiaowei Zhuang. 2021. “Spatially Resolved Cell Atlas of the Mouse Primary Motor Cortex by MERFISH.” Nature 598 (7879): 137–43.

Zhang, Zhuzhu, Jingtian Zhou, Pengcheng Tan, Yan Pang, Angeline C. Rivkin, Megan A. Kirchgessner, Elora Williams, et al. 2021. “Epigenomic Diversity of Cortical Projection Neurons in the Mouse Brain.” Nature 598 (7879): 167–73.

Zingg, Brian, Houri Hintiryan, Lin Gou, Monica Y. Song, Maxwell Bay, Michael S. Bienkowski, Nicholas N. Foster, et al. 2014. “Neural Networks of the Mouse Neocortex.” Cell 156 (5): 1096–1111.

