## Supplementary material for "The BRAIN Initiative Cell Census Network Data Ecosystem: A User’s Guide": BICCN Data Ecosystem - Supplementary

### *Supplementary Materials*

### BICCN Scientific Outcomes

Scientific outcomes of the BICCN have resulted in a dramatically increased understanding of the diversity and consistency of cell types within and between species. A major publication summarizing the initial findings of the consortium for a cross-species analysis of primary motor cortex was published in *Nature,* October 6, 2021, (BRAIN Initiative Cell Census Network (BICCN) 2021). That publication contains 11 publications associated with this consortium that have demonstrated:

- A unified molecular genetic catalog of cortical cell types that integrates transcriptome, open chromatin, and DNA methylation maps (Yao et al. 2021).
- A cross-species analysis that provides a unified taxonomy of transcriptomic types and their hierarchical organization that are conserved from mouse to marmoset and human (BRAIN Initiative Cell Census Network (BICCN) 2021; Bakken et al. 2021).
- Compelling evidence for the epigenomic, transcriptomic, and gene regulatory basis of neuronal phenotypes such as their physiological and anatomical properties, demonstrating the molecular underpinning of neuron types and subtypes (Y. E. Li et al. 2021; Muñoz-Castañeda et al. 2021).
- Identification of spatially resolved cell types through *in situ* single-cell transcriptomics (M. Zhang et al. 2021).
- Development of imaging pipelines and tools for morphometry at whole brain scale, cross modality brain mapping, and resources of single cell morphological cell types (Peng 2021; Qu 2022).
- An extensive genetic toolset for targeting and fate mapping glutamatergic projection neuron types aimed at linking their developmental trajectory to their circuit function (Matho et al. 2021).
- Brain cell type ontologies and nomenclature systems for describing and manipulating cell types (Gillespie et al. 2022; Miller et al. 2020).

Together, the results establish a unified and mechanistic framework of neuronal cell type organization that integrates multimodal molecular, genetic and spatial information with multi-faceted phenotypic properties.

### BICCN Data Levels

A primary value of defining data levels with well-defined structure is in the identification of matching appropriate data with use-cases. The general definition for each level is described below. BICCN workgroups evaluated each of 26 modalities to arrive at a more specific level definition as given in **Suppl. Table 2.**

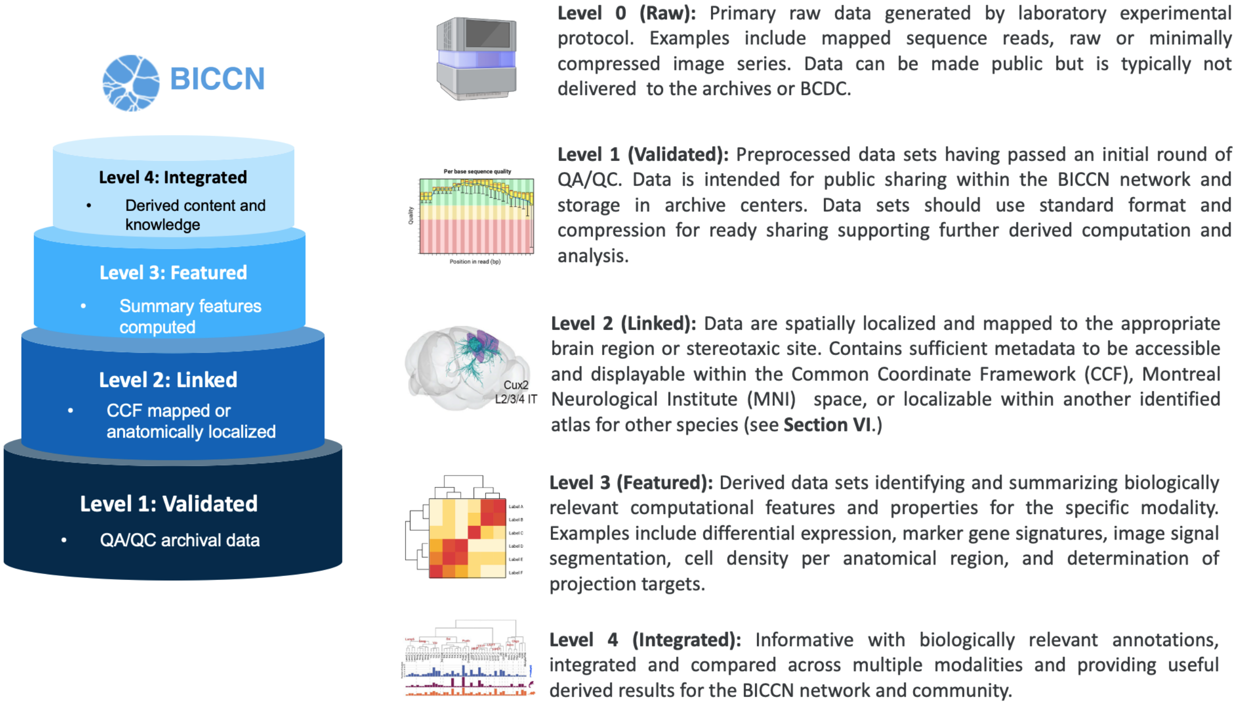


**Supplementary Figure 1: Data Levels.** BICCN data are classified by increasing levels of structure and information content. Level 0 (not shown) raw data, Level 1 QC/QA validated, Level 2 CCF-linked or CCF-mapped, Level 3 feature-derived, Level 4 integrated, annotated. Representative use cases are given below.

- **Level 1 (Validated) Use-Case:** A computational analyst wishes to explore differences in alternative splicing and their significance in variability of cell types, comparing between species and in relationship to the chromatin landscape. **Requirements:** Rigorous QC metrics from transcriptomics and epigenomics modalities. Confidence in mapping alignment is essential (**Level 1**).

- **Level 2 (Linked) Use-Case:** An application developer is building a tool to display long range neuronal connections and cell type specific density distribution in the mouse cortex and corresponding metadata for connection with other modalities. **Requirements:** Reconstructed and QC neurons registered in the CCF, access to mapping transforms for graphical manipulation and display (**Level 2**).

-

- **Level 3 (Featured) Use-Case:** User is working with a Human Lung Cell Atlas (Travaglini et al. 2020; "Human Lung Cell Atlas" n.d.) and wishes to understand differences in cell types found in certain brain regions versus the lung. **Requirements:** This user requires quantified data sets with relevant genomic features identified. Primary interest will be omics-related data sets with spatial regions identified (**Level 3**).

-

- **Level 4 (Integrated) Use-Case:** A software developer in an electron microscopy lab is trying to build an application connecting local neuronal morphology with ultrastructural characteristics. **Requirements:** This user needs spatially resolved local morphology reconstructions with feature quantification in light and EM modalities. (**Levels 3 and 4**)

### Common Coordinate Frameworks

**Mouse CCF Enhancements.** Even for the mouse, one of the most widely used model organisms, the best available histological atlases (e.g., the Allen Reference Atlas (Wang et al. 2020), (Oh et al. 2014)) have significant problems and lack *in vivo* MRI or CT data. Thus, it is not possible to directly determine the coordinate system relevant for stereotactic surgery. Usage of auto fluorescent contrast in place of histochemical contrast has resulted in the loss of cytoarchitectonic information, giving rise to controversies about the location of compartmental boundaries. Further, brainstem annotations and volume are truncated. To address these limitations, we have acquired a multimodal data set, including CT scans, MRI (*in vivo*, *ex vivo*), and histology (Nissl, myelin) data in the same animal. We collect histological series using 3 sectioning planes (coronal, axial, sagittal) at 10-μm section thickness. The sectioning preserves the skull to keep brainstem structures intact. The best existing atlases use one sectioning plane (coronal) and 100-μm spacing. We cross-modally register all datasets and use skull landmarks from *in vivo* MRI and CT to define the stereotactic coordinate system. All datasets are available for viewing and download through our in-house web platform.

**Marmoset CCF.** The new marmoset CCF comprises the averaged *in vivo* T2w MRI volumes of 43 marmosets, (36 female and 7 male), *ex vivo* T2w MRI volumes of 45 marmosets (36 female and 5 male), and high-quality Nissl and myelin histological series from two (1 female and 1 male) subjects. Female and male *in vivo* MRI volumes were also averaged separately, creating CCFs for each sex. The segmentation applied to this CCF interpolates between the RIKEN16 and NIH36 marmoset brain atlases, to fill in the gaps in assigning regional identities to all gray and white matter structures. We use the Paxinos37 print atlas as a supplement to fill in the remaining gaps in segmentation with macroscopic level annotations, and resolve some mismatches across the atlases. The combined NIH36 and RIKEN16 atlas volume with the refined segmentation applied is then registered to the new marmoset CCF.

**Human CCF.** The Brain Architecture Project (http://brainarchitecture.org) is constructing a new CCF for human brains at a previously unprecedented spatial resolution and combining radiological and histological imaging modalities. The new CCF consists of multimodal MRI scans and a series of 3 histological stains (Nissl, myelin, and H&E) carried out on a contiguous set of 20-μm serial sections using the tape-transfer method and reassembled into a 3D multimodal volume. The new CCF rectifies important gaps in previous atlases: The AIBS human reference atlas38 has only 107 histological images. The BigBrain atlas39 from has a complete serial set of approximately 8000 images of 20-μm sections, but only Nissl stained and imaged at low resolution. The new CCF construction pipeline is currently being piloted and refined by two test cases: the creation of a human amygdala and a human hippocampus reference volume.

### Common Coordinate Framework Mapping

Registration techniques and strategy to a CCF are highly dependent on the specific data modality used. Image centric data, where volumetric regions of the brain have been imaged, allows for spatial registration to CCF using more automated image processing tools (**see Section V.2**). These approaches were further developed by the BICCN and use feature detection, image transformation and resampling, and deformable registration methods (Goshtasby 2017; Zitová and Flusser 2003) to obtain quality alignment of source data to the CCFv3.

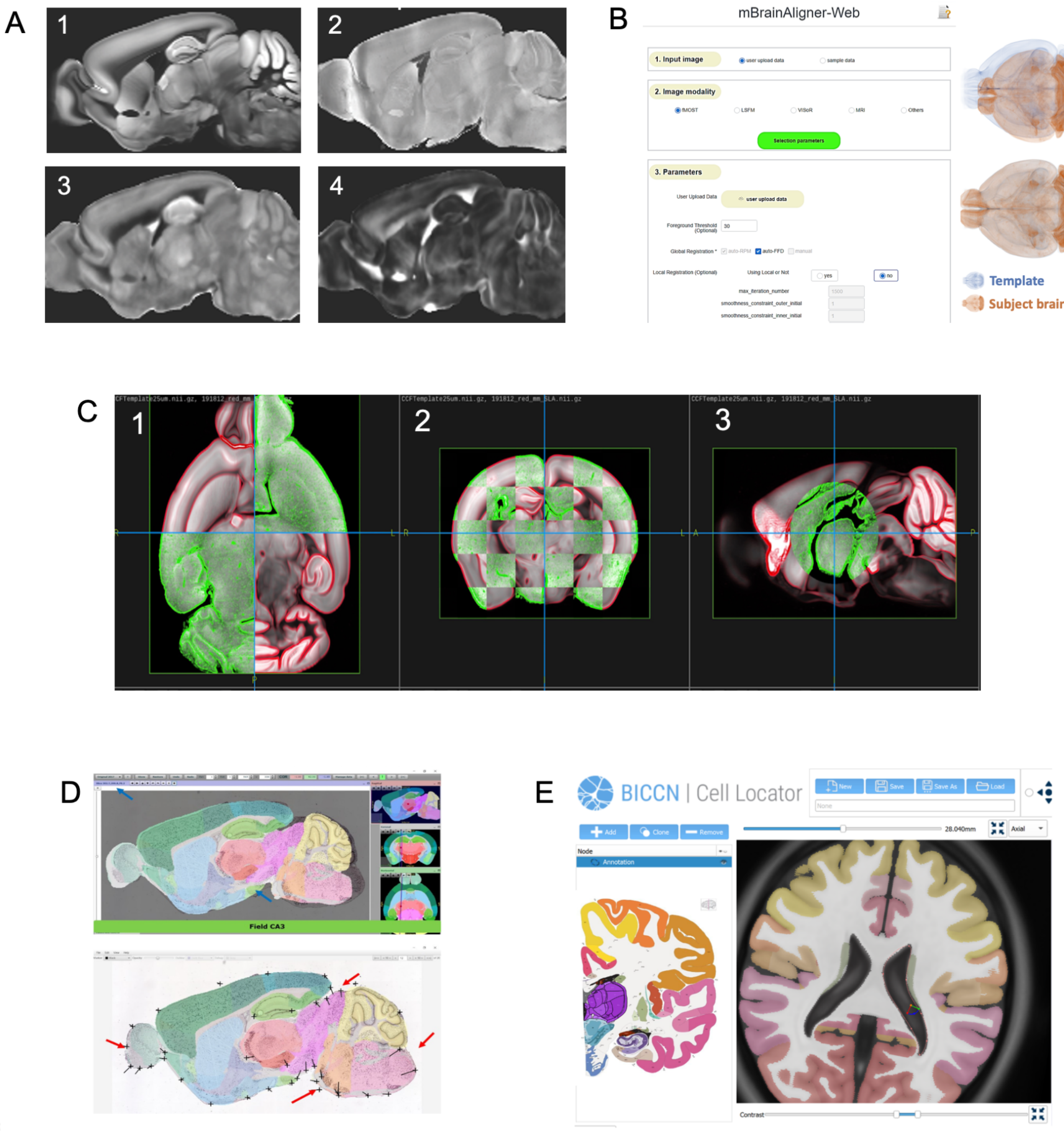


**Supplementary Figure 2. Spatial Registration to Common Coordinate Frameworks.** A) To support image registration to the Allen CCFv3 A1) from different modalities, high resolution templates from MRI and light sheet fluorescence microscopy (LSFM) imaging that are synchronized with CCFv3 have been created (http://dx.doi.org/10.17632/2svx788ddf.1). A2) LSFM based template, A3) MRI and A4) FA contrast based templates. B) Coherent landmark mapping-based registration pipeline mBrainAligner for high-precision cross-modal critical points matching and deformable registration. The tool is available through the web server mBrainAligner-Web. Left panel: The interface of mBrainAligner-Web (http://mbrainaligner.ahu.edu.cn). Right panel: Examples of the subject brain overlaid onto atlas template before a) and after b) registration. C) Software pipeline templates using ANTs (Avants et al. 2008), (Tustison et al. 2021) for deformable registering data from various imaging modalities (e.g., LSFM, MERFISH, fMOST, MRI) were developed and are available via BCDC, as are custom tools, e.g. Entropy, for interactive evaluation and refinement of alignment results, illustrated here in a use case where fMOST data is mapped to CCFv3, C1) horizontal, C2) coronal, C3) sagittal planes. D) EBRAINS tools for registration of 2D images to the CCFv3. Image shown is from the Allen Mouse Brain Atlas, Experiment 75457579. Upper panel: QuickNII (Puchades et al. 2019) is used to register the section to CCFv3 using affine transformations (RRID:SCR_016854) Lower panel: the registration obtained with QuickNII is refined with VisuAlign (RRID:SCR_017978). The atlas is non-linearly deformed over selected target points (black crosses, lower). E) Mapping of human tissue samples is efficiently done using the Cell Locator (https://github.com/BICCN/cell-locator), developed in collaboration with Kitware (www.kitware.com). Based on Slicer technology (www.slicer.org), the human reference annotated atlas serves as data mapping and spatially resolved guides for cell type profiling.

**CCF_Streamlines Package.** A Python package ccf_streamlines (RRID:SCR_022910) was developed to map and visualize the isocortical streamlines of the Allen Mouse Common Coordinate Framework (CCF). These streamlines can be used to visualize three-dimensional CCF-aligned data from the mouse isocortex in flattened or two-dimensional representations, as well as to align data in a consistent pia/white matter orientation. The package allows projecting 3D CCF-aligned data to two-dimensional views and to a 3D slab. The package documentation can be found at https://ccf-streamlines.readthedocs.io/en/latest/index.html.

**Generative Diffeomorphic Mapping for image registration and atlas mapping from the Brain Architecture Portal**. For registration and atlas mapping of multimodal imaging datasets (e.g., datasets combining *in* and *ex vivo* imaging and histology in the same animal/subject), the GDM (Generative Diffeomorphic Mapping) registration algorithm can be employed. Briefly, tissue processing procedures such as extraction and fixation cause brain tissue deformation (Leibnitz 1971; Mouritzen Dam 1979; Schulz et al. 2011) and unguided reconstruction of serial sections leads to accumulated long-range distortions (Malandain et al. 2004). Diffeomorphic mapping emerged to overcome these challenges (Amit et al. 1990; Bajcsy, Lieberson, and Reivich 1983). Our approach to atlas mapping and registration is a generative probabilistic model, where a synthetic stack of 2D microscopy images is formed as a sequence of transforms of a 3D image, plus a noise model describing variability. Transforms include diffeomorphic spatial warping and contrast changes. Mapping to common coordinates is maximum a posteriori (Schmitz and Hof 2011; Woodward et al. 2022, 2018) estimation, and enables reconstruction of 3D and 2D datasets. We quantify tissue distortion with the derivative of spatial maps (morphometry) (Ashburner and Friston 2000), and account for scale changes when quantifying cell density or number (Schmitz and Hof 2011). We support *in vivo* and *ex vivo* MRI, atlas annotations including independently derived segmentation (Schmitz and Hof 2011; Woodward et al. 2022), and multiple stained sections. This framework enables us to jointly analyze multiple MRI contrasts and histology data, providing ground truth data for evaluation of MR models. MRI-constrained reconstruction developed specifically for the marmoset data in our pipeline shows improved accuracy over the baseline method and reduced deformable metric cost. The local scale change between postmortem MRI volumes and reassembled 3D histological volumes from the tape-transfer method is small (~2-3% medial absolute scale change), less than the pre-mortem to postmortem change (~6-10%, measured using the Jacobian determinant of metric tensor relating the corresponding spaces (Lee et al. 2021)). We are currently building a registration toolbox page into the Brain Architecture portal, which will allow users to register their image series to any of the CCFs available on the Brain Architecture portal using an online interface.

### BICCN Image Processing Pipelines

The Image and Multi-Morphology Pipeline (Y. Li et al. 2022; Peng et al. 2010) accesses raw images from the BIL archive (**Section IV.2**) and implements the full pipeline of conversion, processing, morphometry generation, registration and mapping, release, and analysis. The pipeline is hosted on an open cloud platform that features collaborative processing and synergetic computing among various clients, and web interfaces. All data on the server can be accessed through MorphoHub (Jiang et al. 2022), a petabyte-scale multi-morphometry management system and integrates the three largest whole-brain full morphology datasets (Peng et al. 2021a, [b] 2021) (Peng et al. 2021c), MouseLight (Winnubst et al. 2019), and SEU-Allen Institute of neuron reconstruction of the whole brain (Winnubst et al. 2019; Gao et al. 2022).

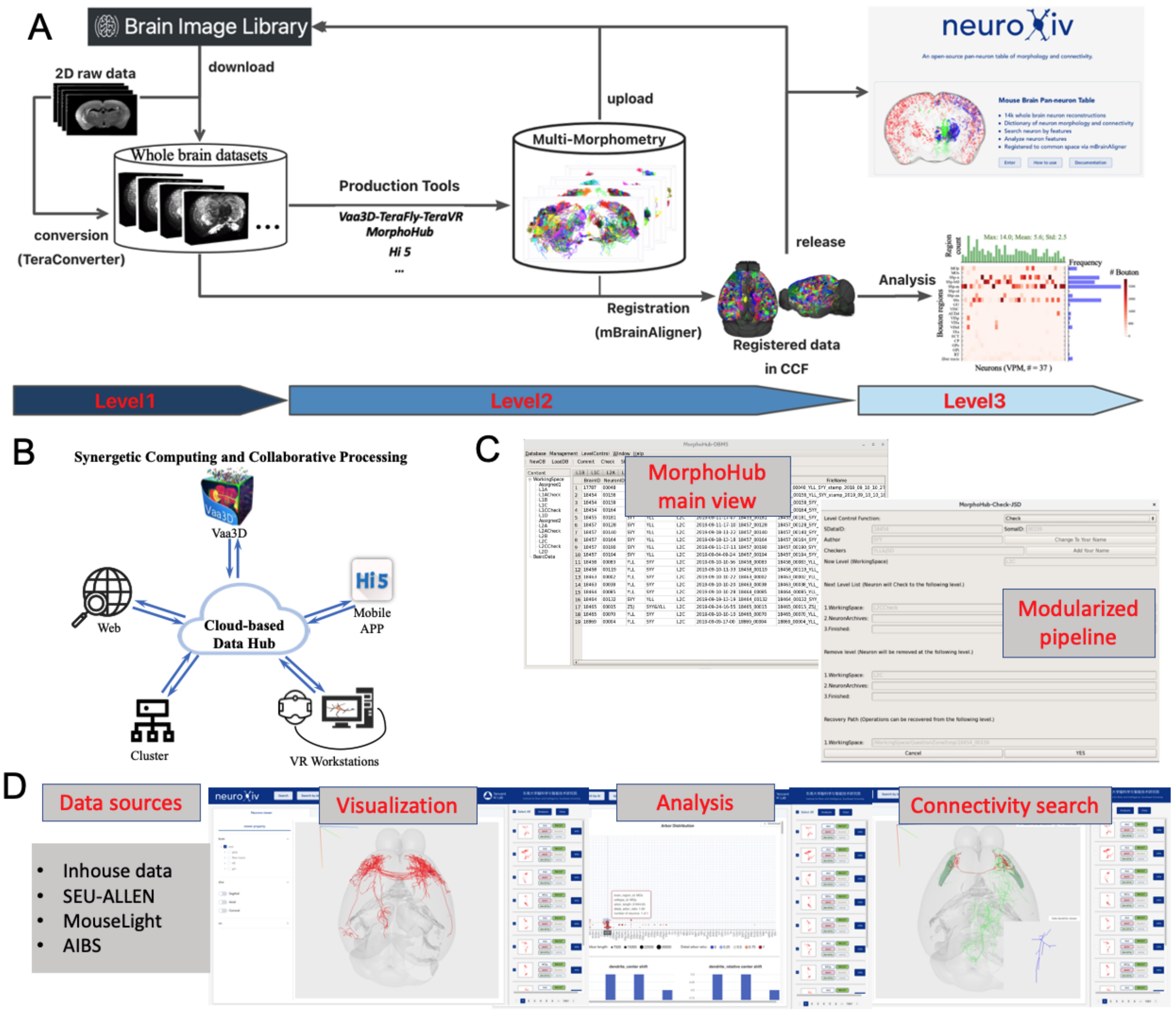


**Supplementary Figure 3. Image and multi-morphology processing pipeline**. A) Schematic pipeline of image processing, multi-morphology production, and mining. The pipeline starts with downloading whole-brain images from public repositories, e.g. BrainImageLibrary, then conversion, processing, morphometry generation, registration and mapping, release, and analysis. B) The pipeline is hosted on an open cloud platform that features collaborative processing and synergetic computing among various clients, including Vaa3D, mobile application Hi5, VR headset system TeraVR, high-performance clusters, and web interfaces, resulting in a dynamic, fast while accurate image and morphometry processing pipeline. C) All data on the server can be accessed through MorphoHub (Jiang et al. 2022), a Petabyte-scale multi-morphometry management system. D) As an open-source pan-neuron table of morphology and connectivity, NeuroXiv integrates the three largest whole-brain full morphology datasets.

### Brain Observatory Storage Service and Database (BossDB, RRID:SCR_017273, https://bossdb.org)

**Brain Observatory Storage Service and Database (BossDB)** is a volumetric, cloud-based data ecosystem for 3D and 4D neuroimaging data (Hider et al. 2022). As the most recent BICCN archive, BossDB focuses primarily on storing volumetric electron microscopy (EM) and X-ray microtomography (XRM) datasets generated as a part of the BRAIN Initiative. BossDB stores high- resolution, multi-channel image data with registered segmentations, annotations, and meshes, and connects to several community resources for data access and data visualization. BossDB also stores connectomics datasets and contains several software tools and interfaces for querying and searching connectomes (Jordan K. Matelsky et al. 2021, 2022). The BossDB ecosystem allows for storing, accessing, and processing multidimensional and volumetric neuroscience datasets through scalable cloud-based resources, and makes use of Amazon Web Services capabilities that ensure available and scalable endpoints, data caching and load balancing, and durable multi-tier data storage. A well-documented interface and API supports a suite of tools, including a Python based software development kit (SDK) that allows a user to easily ingest, validate, visualize, and query neuroscience data from any data generator, making it possible for scientists to discover and share insights on these massive datasets (J. K. Matelsky et al. 2021). For further information see **Suppl. Materials** and BossDB.org | Get Started.

BossDB provides several data ingest paradigms and supports the ingest of several image data formats, where the image and segmentation data is stored in cuboids within a cloud-based object data store. Following data ingestion into BossDB, multiple down-sampled image volume copies are generated and stored to ensure performant data access at multiple image resolutions. BossDB uses a hierarchical organizational structure where it organizes datasets by "collection", "experiment", and "channel". Collections are the top structure containing metadata like the laboratory name and date of creation. Experiments are mid-level structures containing more metadata about the imaging modality, image extent and coordinate frame for the imaged tissue samples (where data exists), resolution levels, and imaging voxel size. Channels, the lowest level structures, contain the volumetric data.

Datasets on the BossDB website (https://bossdb.org/projects) are provided as projects associated with major publications, are publicly available and free to download, and all tools listed are open-source and available for public use (https://bossdb.org/tools). The BossDB provides extensive documentation on the infrastructure, the interfaces, and the various tools that can be used to ingest, access, and visualize volumetric data. A BossDB Cookbook repository (https://github.com/aplbrain/bossdb_cookbook) is a collection of introductory notebooks and examples for interfacing with the BossDB system. For further information see https://bossdb.org/get-started.

### Brain Architecture Portal

All datasets containing high-resolution 2D images from the Mitra Lab1–4, BICCN collaborators (Jeong et al. 2016), and colleagues from other projects (Lin et al. 2019; Majka et al. 2020; Iriki et al. 2018), (Chen et al. 2019; Woodward et al. 2022), (Lin et al. 2019; Majka et al. 2020; Iriki et al. 2018; Okano and Mitra 2015) are available for display on the Brain Architecture web portal. We have extensive experience in serving peta voxels of light microscopic data on the web. The Brain Architecture web portal has been in continuous operation since 2012 and receives 500-1000 unique visitors/month. Datasets are served into species and experiment-type specific pages, accessible from the front landing page (**Suppl. Fig. 4A)**. For example, there is the capability to filter mouse cell distribution datasets via free text search of metadata for keywords (**Suppl. Fig. 4C** and mouse projection and connectivity datasets via injection region or tracer (**Suppl. Fig. 4B**). Clicking on the button on the per dataset metadata card called 'View' or '2D Viewer' will bring up the high-resolution section viewer on the portal (**Suppl. Fig. 4D**). The viewer has in-built capability to display overlays of regional compartments, points indicating cell bodies post cell detection, and skeletons and shaded pixels indicating neuritis post DMM++4 process detection and skeletonization on 2D sections of atlas mapped brains (**Suppl. Fig. 4D**). To date, the viewer displays >2 peta voxels of images using an Angular framework with MySQL DB queries for image file loading directed by Django-based APIs. The viewer can display data at multiple resolutions, with zoom to super resolution capability, beyond the native in-plane 0.46-µm resolution of the images.

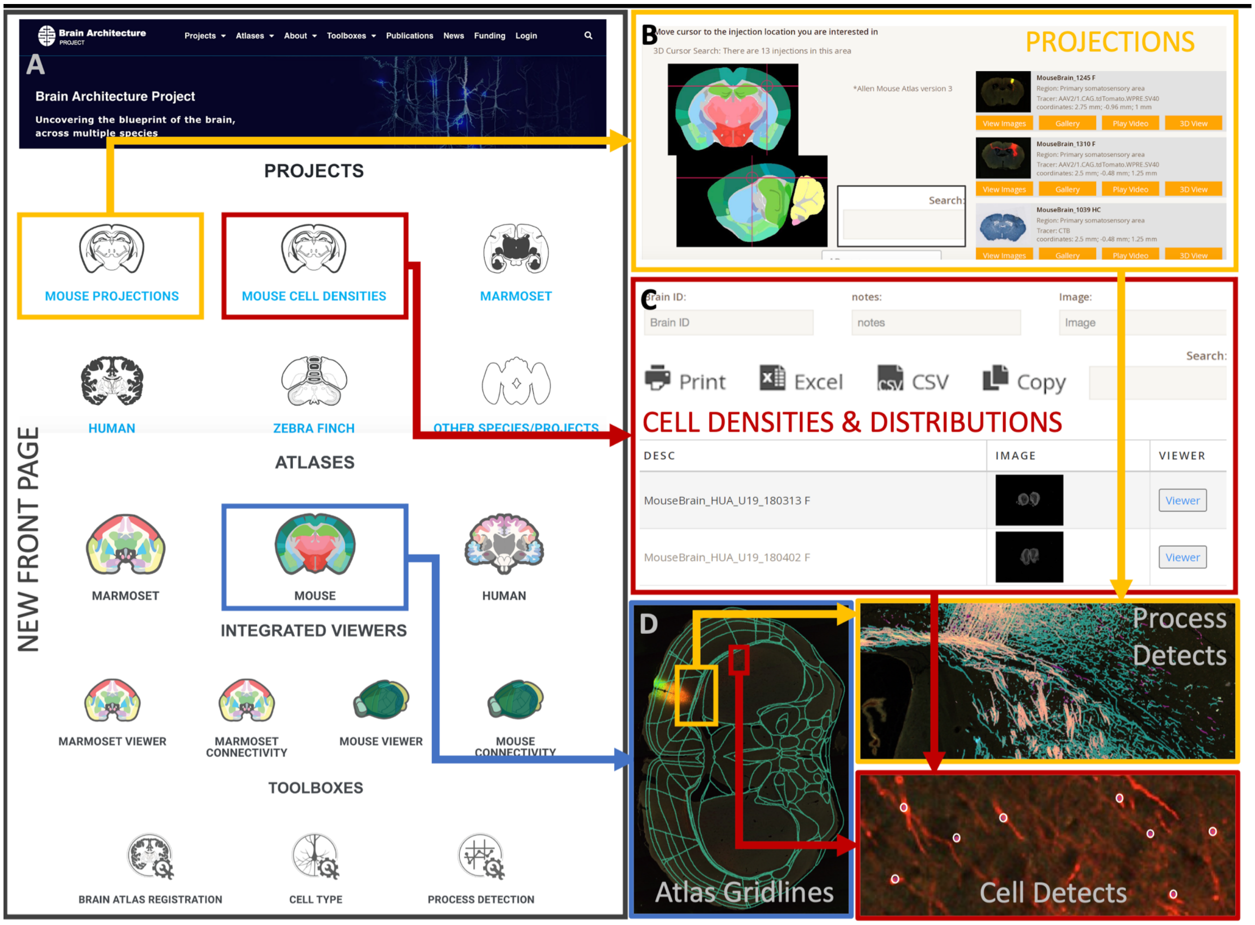

A
Brain Architecture Project
Projects Atlases About Toolboxes Publications News Funding Login
Brain Architecture Project
Uncovering the blueprint of the brain, across multiple species
NEW FRONT PAGE
PROJECTS
MOUSE PROJECTIONS
MOUSE CELL DENSITIES
MARMOSET
HUMAN
ZEBRA FINCH
OTHER SPECIES/PROJECTS
ATLASES
MARMOSET
MOUSE
HUMAN
INTEGRATED VIEWERS
MARMOSET VIEWER
MARMOSET CONNECTIVITY
MOUSE VIEWER
MOUSE CONNECTIVITY
TOOLBOXES
BRAIN ATLAS REGISTRATION
CELL TYPE
PROCESS DETECTION
B
Move cursor to the injection location you are interested in
3D Cursor Search: There are 13 injections in this area
*Allen Mouse Atlas version 3
Search:
PROJECTIONS
MouseBrain_1245 F
MouseBrain_1310 F
MouseBrain_1039 HC
View Images
Gallery
Play Video
3D View
C
Brain ID:
notes:
Image:
Search:
Print
Excel
CSV
Copy
CELL DENSITIES & DISTRIBUTIONS
DESC
IMAGE
VIEWER
MouseBrain_HUA_U19_180313 F
Viewer
MouseBrain_HUA_U19_180402 F
Viewer
D
Atlas Gridlines
Process Detects
Cell Detects

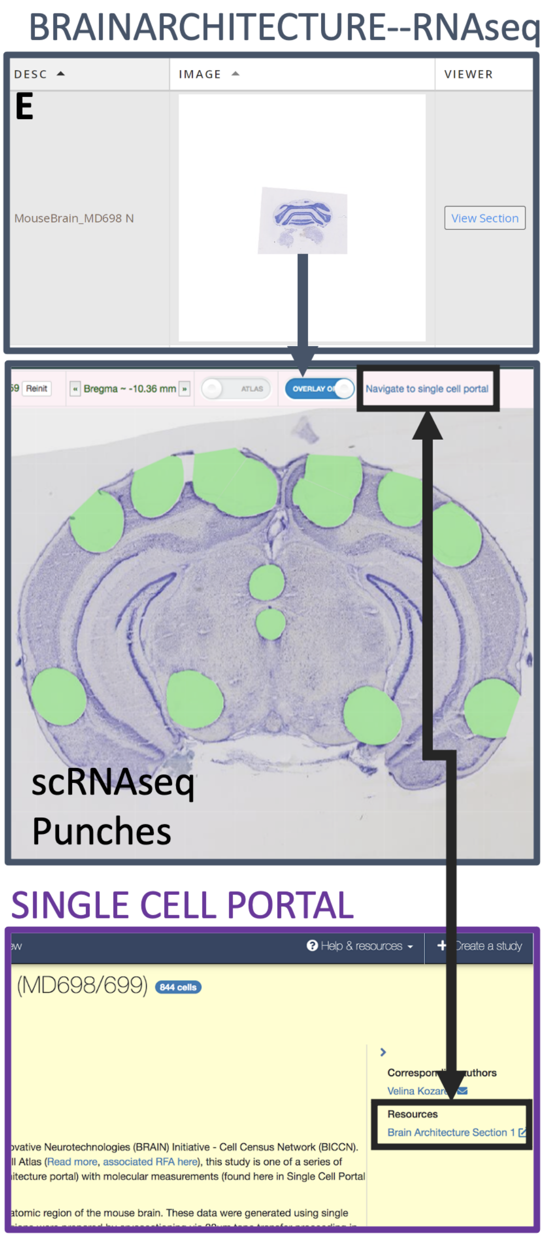


**Supplementary Figure 4**. A graphical description of the data types available and use of the Brain Architecture portal and high-resolution 2D viewer. A) An image of the new front page. Clicking on the buttons will lead to subsidiary data and atlas selection pages. B) Following the gold boxes and arrows, clicking on the Mouse Projections page will lead to a dataset selection page as pictured, while C) following the red boxes and arrows, clicking on the Mouse Cell Densities page will lead to a different selection page. D) Clicking on the 'Viewer' button on any dataset selection page, or selecting a 2D view of an atlas, will lead to the high-resolution 2D viewer. Atlas 2D views will contain regional gridlines post atlas mapping and registration. Process detections and skeletons from projection data and cell body or nucleus detections from cell distribution data can also be overlaid on the 2D images, as pictured. E) Established cross-linkage between the Brain Architecture portal and the Single-Cell Portal from Broad Institute, as indicated by the arrows.

In addition to thousands of viewable datasets across multiple projects, all software tool sets employed in projects involving the Mitra laboratory image analytics pipeline, including GDM registration and atlas mapping (Tward et al., n.d.), cell detection3, and process detection and skeletonization via DMM++4, will be available both in interactive versions on the Brain Architecture web portal, and for download (of both source code and documentation) on Github and Bitbucket repositories; buttons to access these can be found on the new front page (**Suppl. Fig. 4A**). Interactive analytic tool sets will have their own dedicated pages on the Brain Architecture web portal and will be open to use for any user who creates a free account. All downloads from Github and Bitbucket repositories will be unrestricted, in keeping with Open Source code practices.

### BICCN Tools and Resources

Many essential tools and resources were developed throughout the BICCN and are summarized below and on the portal https://biccn.org/tools.

- **Epiviz** (RRID:SCR_022796**),** an interactive visualization tool for functional genomics data; *Brainome* (https://brainome.ucsd.edu/annoj/BICCN_MOp/) a genome browser to visualize the cell type-specific transcriptomes and epigenomes of cell types from the mouse MOp.
- **Catlas** (RRID:SCR_018690), which provides maps of accessible chromatin in >800,000 individual cells from 45 regions spanning the adult mouse isocortex, olfactory bulb, hippocampus, and cerebral nuclei.
- C**han Zuckerberg Initiative (CZI, RRID:SCR_021059)** CZ CELL x GENE is a web-based interface for exploring high dimensional datasets along categorical, continuous and spatial dimensions, as well as feature annotation and hosts several of the molecular datasets of the BICCN, Human Cell Atlas (https://www.humancellatlas.org, HCA, RRID:SCR_016530), and and other consortia including an atlas of cortical arealization in developing human CZ CELL x GENE (Bhaduri et al. 2021).
- **Cytosplore Viewer** (RRID:SCR_018330 is a stand-alone application (Windows and MacOS) for interactive visual exploration of multi-species and cross-omics single cell data in several BICCN

data resources. It supports interactive exploration of cellular hierarchies, gene expression and metadata of individual cells, and it allows computation of differential statistics between cell selections or clusters, and across species. Cytosplore Viewer enables linked cross-omics, cross-species comparison between matched gene expression, chromatin state and DNA methylation data through linkout to the UCSC genome browser.

- **CloudVolume** (RRID:SCR_022820) is a Python package that allows for easy reading and writing of image data in the neuroglancer precomputed format (https://github.com/google/neuroglancer/tree/master/src/neuroglancer/datasource/precomputed). The precomputed format allows for efficient visualization of large 3D images by exploiting the fact that at any given moment, users typically only need to view a small subset of an image, i.e. a certain spatial area at a certain resolution. Many of the datasets hosted at BossDB (https://bossdb.org/projects) are available in the precomputed format, and therefore the raw digital data can be read in Python using CloudVolume.
- **MetaNeighbor** (Crow et al. 2018; Fischer et al. 2021)RRID:SCR_016727, (Crow et al. 2018; Fischer et al. 2021) is a method for assessing the replicability of single cell data, used in a number of key BICCN publications (e.g., (Bakken et al. 2021; Yao et al. 2021)) to validate cell types and perform quality control. Rather than merging data, MetaNeighbor holds data independent and tests the degree to which cell-types in one dataset can be used to characterize the cell-types in another dataset. This highly scalable procedure can be easily applied to test the specific replicability of particular genes sets. Because the method does not modify the underlying data, it is useful for comparisons where strong real potential differences exist (e.g., cross-species assessment). The R version of MetaNeighbor is available through Bioconductor at https://www.bioconductor.org/packages/release/bioc/html/MetaNeighbor.html. The development versions are available on GitHub at https://github.com/gillislab/MetaNeighbor (R version) and https://github.com/gillislab/pyMN (Python version). The related method MetaMarkers, provides meta-analytic marker sets across the BICCN data corpus (Fischer and Gillis 2021).
- **NS-Forest** (RRID:SCR_018348) is a Python package that identifies minimum sets of marker genes for cell types identified in sc/snRNA-seq datasets (B. D. Aevermann et al. 2018; B. Aevermann et

al. 2021). The method is based on the selection of informative gene expression features using random forest machine learning and includes a binary scoring step to enrich for genes that show binary on-off expression patterns, which are especially useful for downstream applications like spatial transcriptomics. NS-Forest markers have been used to annotate cell types identified in BICCN datasets (Bakken et al. 2021) and form the basis of cell type definition in the Brain Data Standards Ontology described below and for visualization in the CTKE.

- **FR-Match** (https://github.com/JCVenterInstitute/FRmatch) is an R package for matching cell types across datasets based on a novel application of the Friedman-Rafsky (FR) test, a non-parametric statistical test for multivariate data comparison, in the context of single cell clustering results. (Y. Zhang et al. 2021). FR-Match has been validated for matching single nucleus and single cell datasets, matching data from SmartSeq and 10X Genomics platforms, and matching single cell and spatial transcriptomics datasets using BICCN datasets (Y. Zhang et al. 2022), illustrating the value of the BICCN data ecosystem for novel methods development.
- **Single Cell Portal (**RRID:SCR_014816) is a portal to visualize, share, and disseminate single-cell data. As a self-service application, scientists upload data programmatically or use a wizard-driven UI with the help of extensive support (help desk, documentation wiki, and API swagger). Studies allow for a collection of rich interactive visualizations and customizations and are findable through natural language search or curated search facets. Single Cell Portal includes an end-to-end set of functionality including private, sharing, reviewer, and public settings. This portal is generally available to the scientific community and houses collections associated with the BICCN, the Human Cell Atlas project, the Alexandria project, and COVID-19 research.

### BICCN FAIR Data Practices

The **Findability** of pipelines is enabled through the use of RRIDs to give pipelines unique, explicit identifiers. These pipelines are under continuous development and so semantic versioning is used to refer to versions and specify a specific commit of the code. The BICCN Portal hosts a page that collects links to all resources (code repositories, Terra workspaces, publications, and documentation) in one place. To assure **Accessibility**, the BCDC hosts pipelines in multiple community resources including: public GitHub repositories (for software engineers), Dockstore (for computational biologists), and Terra (installed and ready to run for those without local infrastructure or who want to use scalable cloud resources). Pipelines are open-access and freely licensed to encourage not-for-profit as well as for-profit use. All pipelines are fully documented and are preinstalled into Terra with test data and expected outputs to remove the requirement of complicated installations or infrastructure requirements. **Interoperability** for pipelines is addressed for both infrastructures to run pipelines as well as interoperability with other tools or pipelines. Execution interoperability is encouraged by using a modern work flow language (WDL) that separates the code performing scientific tasks from code for orchestrating the pipeline on infrastructure (contained in workflow execution programs). WDL can be executed by multiple workflow execution programs including **Cromwell,** a portable execution engine leveraged in high-throughput sequencing settings that can be launched in many environments. To encourage interoperability with other tools and pipelines, pipelines leverage *de facto* and GA4GH (www.ga4gh.org) standards and preference leveraging community developed tools over creating new tools. Finally, all these activities come together to enable essential **Reproducible** science.

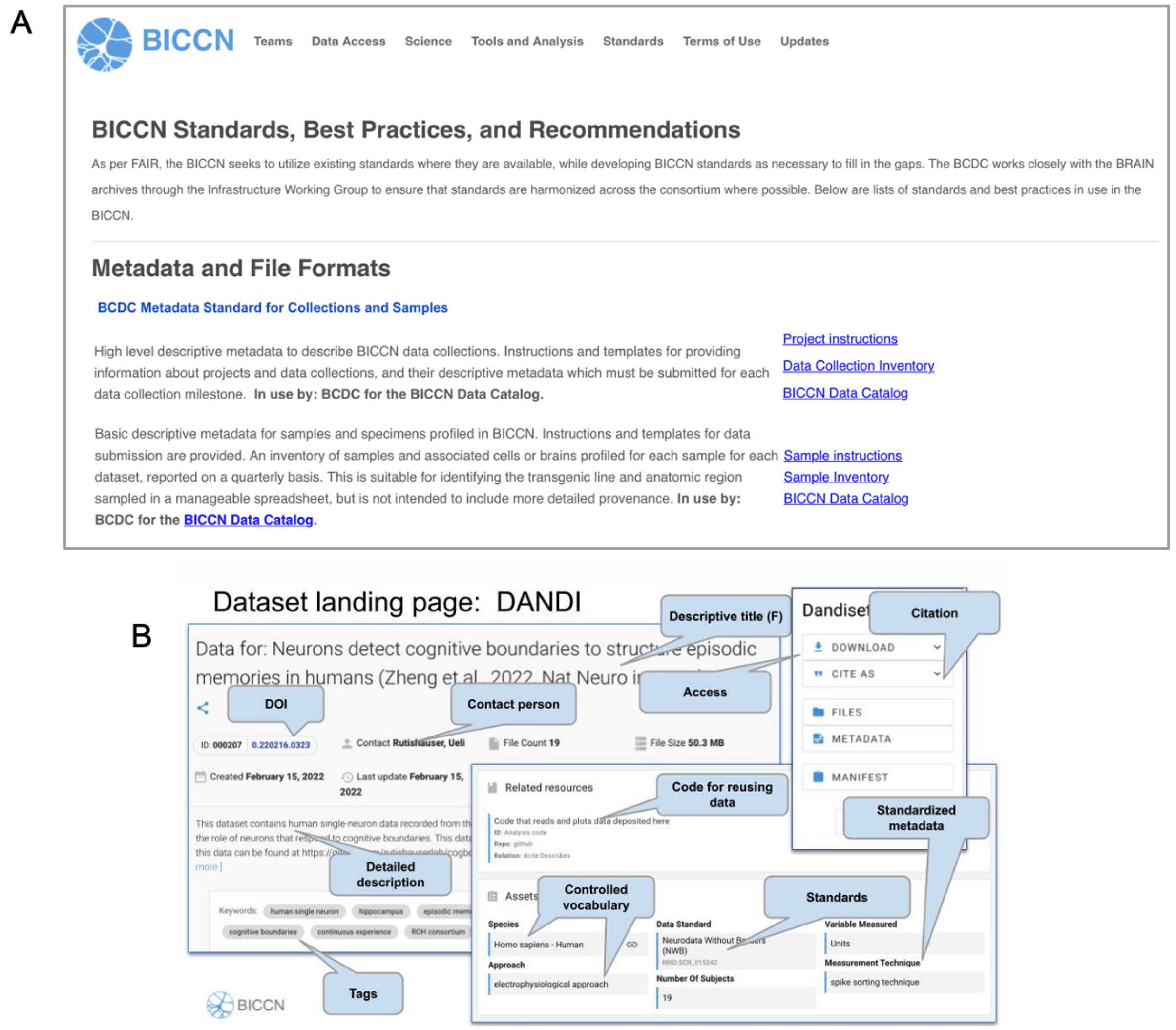


**Figure 5. DANDI Archive and FAIR data practices**. Annotated dataset landing page from the DANDI archive illustrating key characteristics of FAIR data description including DOI, reference to contact individual, details description of the data set with metadata and controlled vocabulary. Reference to utilized standards, access and code for reusing the dataset are provided. All Dandisets have similar consistent documentation.

### Brain Data Standards Ontology

The BDSO approach is described in detail in Tan et al. {(Tan et al., n.d.)), not only makes it scalable, but also lowers human error (as compared to manually creating the ontology). These features are crucial as we scale up to whole brains in the new phase described in the next section. BDSO is developed based on the OBO Foundry (Jackson et al. 2021) and FAIR (Jackson et al. 2021; Wilkinson et al. 2016) principles; as such, the BDSO is fully compliant with OBO Foundry standards and has been included as an OBO Foundry ontology. The methods we described for creating BDSO were made to be relatively generalizable, allowing future projects to adapt them with BDSO serving as an example (Tan et al., n.d.).

In their native formats, ontologies are not practical for driving web applications. The BDSO loads into an instance of the graph database Neo4j using a standard pipeline for logical projection of OWL into a Neo4j property graph {https://doi.org/10.5281/zenodo.7082530}, using standard ontology queries to add tags to drive faceted searching for neurons by type (GABAergic), morphology (Chandelier) location (Primary Motor Cortex) and species. The structure of the knowledge graph is used to populate and index documents in a SOLR document store, enabling enhanced full-text search and content retrieval for the CTKE.

The BDSO's code base is available at GitHub (RRID:SCR_022822) including documentation of the full technology stack and details of the approach. The indexer, indexes and API code for the CTKE are available at ("[No Title]" n.d.) The latest release of the ontology is available for download from http://purl.obolibrary.org/obo/pcl/bds/bds.owl and is hosted on the EMBL-EBI ontology lookup service OLS (http://ceur-ws.org/Vol-1546/paper_29.pdf) at https://www.ebi.ac.uk/ols/ontologies/pcl. OLS provides ontology search, browsing, visualization capabilities and enables web services driven programmatic access to the BDSO.

Aevermann, Brian D., Mark Novotny, Trygve Bakken, Jeremy A. Miller, Alexander D. Diehl, David Osumi-Sutherland, Roger S. Lasken, Ed S. Lein, and Richard H. Scheuermann. 2018. "Cell Type Discovery Using Single-Cell Transcriptomics: Implications for Ontological Representation." *Human Molecular Genetics* 27 (R1): R40–47.

Aevermann, Brian, Yun Zhang, Mark Novotny, Mohamed Keshk, Trygve Bakken, Jeremy Miller, Rebecca Hodge, Boudewijn Lelieveldt, Ed Lein, and Richard H. Scheuermann. 2021. "A Machine Learning Method for the Discovery of Minimum Marker Gene Combinations for Cell Type Identification from Single-Cell RNA Sequencing." *Genome Research* 31 (10): 1767–80.

Amit, Y., Ulf Grenander, M. Piccioni, and Center for Intelligent Control Systems (U.S.). 1990. *Structural Image Restoration Through Deformable Templates*.

Ashburner, J., and K. J. Friston. 2000. "Voxel-Based Morphometry--the Methods." *NeuroImage* 11 (6 Pt 1): 805–21.

Avants, B. B., C. L. Epstein, M. Grossman, and J. C. Gee. 2008. "Symmetric Diffeomorphic Image Registration with Cross-Correlation: Evaluating Automated Labeling of Elderly and

Neurodegenerative Brain.” *Medical Image Analysis* 12 (1): 26–41.
Bajcsy, R., R. Lieberson, and M. Reivich. 1983. “A Computerized System for the Elastic Matching of Deformed Radiographic Images to Idealized Atlas Images.” *Journal of Computer Assisted Tomography* 7 (4): 618–25.
Bakken, Trygve E., Nikolas L. Jorstad, Qiwen Hu, Blue B. Lake, Wei Tian, Brian E. Kalmbach, Megan Crow, et al. 2021. “Comparative Cellular Analysis of Motor Cortex in Human, Marmoset and Mouse.” *Nature* 598 (7879): 111–19.
Bhaduri, Aparna, Carmen Sandoval-Espinosa, Marcos Otero-Garcia, Irene Oh, Raymund Yin, Ugomma C. Eze, Tomasz J. Nowakowski, and Arnold R. Kriegstein. 2021. “An Atlas of Cortical Arealization Identifies Dynamic Molecular Signatures.” *Nature* 598 (7879): 200–204.
BRAIN Initiative Cell Census Network (BICCN). 2021. “A Multimodal Cell Census and Atlas of the Mammalian Primary Motor Cortex.” *Nature* 598 (7879): 86–102.
Chen, Yuncong, Lauren E. McElvain, Alexander S. Tolpygo, Daniel Ferrante, Beth Friedman, Partha P. Mitra, Harvey J. Karten, Yoav Freund, and David Kleinfeld. 2019. “An Active Texture-Based Digital Atlas Enables Automated Mapping of Structures and Markers across Brains.” *Nature Methods* 16 (4): 341–50.
Crow, Megan, Anirban Paul, Sara Ballouz, Z. Josh Huang, and Jesse Gillis. 2018. “Characterizing the Replicability of Cell Types Defined by Single Cell RNA-Sequencing Data Using MetaNeighbor.” *Nature Communications* 9 (1): 1–12.
Fischer, Stephan, Megan Crow, Benjamin D. Harris, and Jesse Gillis. 2021. “Scaling up Reproducible Research for Single-Cell Transcriptomics Using MetaNeighbor.” *Nature Protocols* 16 (8): 4031–67.
Fischer, Stephan, and Jesse Gillis. 2021. “How Many Markers Are Needed to Robustly Determine a Cell’s Type?” *iScience* 24 (11): 103292.
Gao, Le, Sang Liu, Lingfeng Gou, Yachuang Hu, Yanhe Liu, Li Deng, Danyi Ma, et al. 2022. “Single-Neuron Projectome of Mouse Prefrontal Cortex.” *Nature Neuroscience* 25 (4): 515–29.
Gillespie, Thomas H., Shreejoy J. Tripathy, Mohameth François Sy, Maryann E. Martone, and Sean L. Hill. 2022. “The Neuron Phenotype Ontology: A FAIR Approach to Proposing and Classifying Neuronal Types.” *Neuroinformatics*, March. https://doi.org/10.1007/s12021-022-09566-7.
Goshtasby, Arthur Ardeshir. 2017. *Theory and Applications of Image Registration*. John Wiley & Sons.
Hider, Robert, Jr, Dean Kleissas, Timothy Gion, Daniel Xenes, Jordan Matelsky, Derek Pryor, Luis Rodriguez, Erik C. Johnson, William Gray-Roncal, and Brock Wester. 2022. “The Brain Observatory Storage Service and Database (BossDB): A Cloud-Native Approach for Petascale Neuroscience Discovery.” *Frontiers in Neuroinformatics* 0. https://doi.org/10.3389/fninf.2022.828787.
“Human Lung Cell Atlas.” n.d. Accessed October 23, 2022. https://hlca.ds.czbiohub.org.
Iriki, Atsushi, Hirotaka James Okano, Erika Sasaki, and Hideyuki Okano. 2018. *The 3-Dimensional Atlas of the Marmoset Brain: Reconstructible in Stereotaxic Coordinates*. Springer.
Jackson, Rebecca, Nicolas Matentzoglu, James A. Overton, Randi Vita, James P. Balhoff, Pier Luigi Buttigieg, Seth Carbon, et al. 2021. “OBO Foundry in 2021: Operationalizing Open Data Principles to Evaluate Ontologies.” *Database: The Journal of Biological Databases and Curation* 2021 (October). https://doi.org/10.1093/database/baab069.
Jeong, Minju, Yongsoo Kim, Jeongjin Kim, Daniel D. Ferrante, Partha P. Mitra, Pavel Osten, and Daesoo Kim. 2016. “Comparative Three-Dimensional Connectome Map of Motor Cortical Projections in the Mouse Brain.” *Scientific Reports* 6 (February): 20072.
Jiang, Shengdian, Yimin Wang, Lijuan Liu, Liya Ding, Zongcai Ruan, Hong-Wei Dong, Giorgio A. Ascoli, Michael Hawrylycz, Hongkui Zeng, and Hanchuan Peng. 2022. “Petabyte-Scale Multi-Morphometry of Single Neurons for Whole Brains.” *Neuroinformatics*, February. https://doi.org/10.1007/s12021-022-09569-4.
Lee, Brian C., Meng K. Lin, Yan Fu, Junichi Hata, Michael I. Miller, and Partha P. Mitra. 2021. “Multimodal

Cross-Registration and Quantification of Metric Distortions in Marmoset Whole Brain Histology Using Diffeomorphic Mappings." *The Journal of Comparative Neurology* 529 (2): 281–95.

Leibnitz, L. 1971. "[Studies on the optimation of weight and volume changes in brains during fixation, dehydration and brightening as well as on fresh brain volume estimates based on the weight of treated brains]." *Journal fur Hirnforschung* 13 (4): 320–29.

Lin, Meng Kuan, Yeonsook Shin Takahashi, Bing-Xing Huo, Mitsutoshi Hanada, Jaimi Nagashima, Junichi Hata, Alexander S. Tolpygo, et al. 2019. "A High-Throughput Neurohistological Pipeline for Brain-Wide Mesoscale Connectivity Mapping of the Common Marmoset." *eLife* 8 (February). https://doi.org/10.7554/eLife.40042.

Li, Yang Eric, Sebastian Preissl, Xiaomeng Hou, Ziyang Zhang, Kai Zhang, Yunjiang Qiu, Olivier B. Poirion, et al. 2021. "An Atlas of Gene Regulatory Elements in Adult Mouse Cerebrum." *Nature* 598 (7879): 129–36.

Li, Yuanyuan, Jun Wu, Donghuan Lu, Chao Xu, Yefeng Zheng, Hanchuan Peng, and Lei Qu. 2022. "mBrainAligner-Web: A Web Server for Cross-Modal Coherent Registration of Whole Mouse Brains." *Bioinformatics* , August. https://doi.org/10.1093/bioinformatics/btac549.

Majka, Piotr, Shi Bai, Sophia Bakola, Sylwia Bednarek, Jonathan M. Chan, Natalia Jermakow, Lauretta Passarelli, et al. 2020. "Open Access Resource for Cellular-Resolution Analyses of Corticocortical Connectivity in the Marmoset Monkey." *Nature Communications* 11 (1): 1133.

Malandain, Grégoire, Éric Bardinet, Koen Nelissen, and Wim Vanduffel. 2004. "Fusion of Autoradiographs with an MR Volume Using 2-D and 3-D Linear Transformations." *NeuroImage*. https://doi.org/10.1016/j.neuroimage.2004.04.038.

Matelsky, J. K., L. M. Rodriguez, D. Xenes, T. Gion, R. Hider, B. A. Wester, and W. Gray-Roncal. 2021. "An Integrated Toolkit for Extensible and Reproducible Neuroscience." *Conference Proceedings: ... Annual International Conference of the IEEE Engineering in Medicine and Biology Society. IEEE Engineering in Medicine and Biology Society. Conference* 2021 (November). https://doi.org/10.1109/EMBC46164.2021.9630199.

Matelsky, Jordan K., Erik C. Johnson, Brock Wester, and William Gray-Roncal. 2022. "Scalable Graph Analysis Tools for the Connectomics Community." *bioRxiv*. https://doi.org/10.1101/2022.06.01.494307.

Matelsky, Jordan K., Elizabeth P. Reilly, Erik C. Johnson, Jennifer Stiso, Danielle S. Bassett, Brock A. Wester, and William Gray-Roncal. 2021. "DotMotif: An Open-Source Tool for Connectome Subgraph Isomorphism Search and Graph Queries." *Scientific Reports* 11 (1): 1–14.

Matho, Katherine S., Dhananjay Huilgol, William Galbavy, Miao He, Gukhan Kim, Xu An, Jiangteng Lu, et al. 2021. "Genetic Dissection of the Glutamatergic Neuron System in Cerebral Cortex." *Nature*. https://doi.org/10.1038/s41586-021-03955-9.

Miller, Jeremy A., Nathan W. Gouwens, Bosiljka Tasic, Forrest Collman, Cindy Tj van Velthoven, Trygve E. Bakken, Michael J. Hawrylycz, Hongkui Zeng, Ed S. Lein, and Amy Bernard. 2020. "Common Cell Type Nomenclature for the Mammalian Brain." *eLife* 9 (December). https://doi.org/10.7554/eLife.59928.

Mouritzen Dam, A. 1979. "Shrinkage of the Brain during Histological Procedures with Fixation in Formaldehyde Solutions of Different Concentrations." *Journal Fur Hirnforschung* 20 (2): 115–19.

Muñoz-Castañeda, Rodrigo, Brian Zingg, Katherine S. Matho, Xiaoyin Chen, Quanxin Wang, Nicholas N. Foster, Anan Li, et al. 2021. "Cellular Anatomy of the Mouse Primary Motor Cortex." *Nature* 598 (7879): 159–66.

"[No Title]." n.d. Accessed September 24, 2022. http://ceur-ws.org/Vol-1546/paper_29.pdf.

Oh, Seung Wook, Julie A. Harris, Lydia Ng, Brent Winslow, Nicholas Cain, Stefan Mihalas, Quanxin Wang, et al. 2014. "A Mesoscale Connectome of the Mouse Brain." *Nature* 508 (7495): 207–14.

Okano, Hideyuki, and Partha Mitra. 2015. "Brain-Mapping Projects Using the Common Marmoset."

*Neuroscience Research* 93 (April): 3–7.
Peng, Hanchuan, Zongcai Ruan, Fuhui Long, Julie H. Simpson, and Eugene W. Myers. 2010. "V3D Enables Real-Time 3D Visualization and Quantitative Analysis of Large-Scale Biological Image Data Sets." *Nature Biotechnology* 28 (4): 348–53.
Peng, Hanchuan, Peng Xie, Lijuan Liu, Xiuli Kuang, Yimin Wang, Lei Qu, Hui Gong, et al. 2021a. "Morphological Diversity of Single Neurons in Molecularly Defined Cell Types." *Nature* 598 (7879): 174–81.
———. 2021b. "Morphological Diversity of Single Neurons in Molecularly Defined Cell Types." *Nature* 598 (7879): 174–81.
———. 2021c. "Morphological Diversity of Single Neurons in Molecularly Defined Cell Types." *Nature* 598 (7879): 174–81.
Puchades, Maja A., Gergely Csucs, Debora Ledergerber, Trygve B. Leergaard, and Jan G. Bjaalie. 2019. "Spatial Registration of Serial Microscopic Brain Images to Three-Dimensional Reference Atlases with the QuickNII Tool." *PloS One* 14 (5): e0216796.
Schmitz, Christoph, and Patrick R. Hof. 2011. "Design-Based Stereology in Brain Aging Research." In *Brain Aging: Models, Methods, and Mechanisms*, edited by David R. Riddle. Boca Raton (FL): CRC Press/Taylor & Francis.
Schulz, Georg, Hendrikus J. A. Crooijmans, Marco Germann, Klaus Scheffler, Magdalena Müller-Gerbl, and Bert Müller. 2011. "Three-Dimensional Strain Fields in Human Brain Resulting from Formalin Fixation." *Journal of Neuroscience Methods* 202 (1): 17–27.
Tan, Shawn Zheng Kai, Huseyin Kir, Brian Aevermann, Tom Gillespie, Michael Hawrylycz, Ed Lein, Nicolas Matentzoglu, et al. n.d. "Brain Data Standards Ontology: A Data-Driven Ontology of Transcriptomically Defined Cell Types in the Primary Motor Cortex." https://doi.org/10.1101/2021.10.10.463703.
Travaglini, Kyle J., Ahmad N. Nabhan, Lolita Penland, Rahul Sinha, Astrid Gillich, Rene V. Sit, Stephen Chang, et al. 2020. "A Molecular Cell Atlas of the Human Lung from Single-Cell RNA Sequencing." *Nature* 587 (7835): 619–25.
Tustison, Nicholas J., Philip A. Cook, Andrew J. Holbrook, Hans J. Johnson, John Muschelli, Gabriel A. Devenyi, Jeffrey T. Duda, et al. 2021. "The ANTsX Ecosystem for Quantitative Biological and Medical Imaging." *Scientific Reports* 11 (1): 9068.
Tward, Daniel, Xu Li, Bingxing Huo, Brian Lee, Michael I. Miller, and Partha P. Mitra. n.d. "Solving the *where* Problem in Neuroanatomy: A Generative Framework with Learned Mappings to Register Multimodal, Incomplete Data into a Reference Brain." https://doi.org/10.1101/2020.03.22.002618.
Wang, Quanxin, Song-Lin Ding, Yang Li, Josh Royall, David Feng, Phil Lesnar, Nile Graddis, et al. 2020. "The Allen Mouse Brain Common Coordinate Framework: A 3D Reference Atlas." *Cell* 181 (4): 936–53.e20.
Wilkinson, Mark D., Michel Dumontier, I. Jsbrand Jan Aalbersberg, Gabrielle Appleton, Myles Axton, Arie Baak, Niklas Blomberg, et al. 2016. "The FAIR Guiding Principles for Scientific Data Management and Stewardship." *Scientific Data* 3 (March): 160018.
Winnubst, Johan, Erhan Bas, Tiago A. Ferreira, Zhuhao Wu, Michael N. Economo, Patrick Edson, Ben J. Arthur, et al. 2019. "Reconstruction of 1,000 Projection Neurons Reveals New Cell Types and Organization of Long-Range Connectivity in the Mouse Brain." *Cell* 179 (1): 268–81.e13.
Woodward, Alexander, Tsutomu Hashikawa, Masahide Maeda, Takaaki Kaneko, Keigo Hikishima, Atsushi Iriki, Hideyuki Okano, and Yoko Yamaguchi. 2018. "The Brain/MINDS 3D Digital Marmoset Brain Atlas." *Scientific Data* 5 (February): 180009.
———. 2022. "Author Correction: The Brain/MINDS 3D Digital Marmoset Brain Atlas." *Scientific Data* 9 (1): 100.
Yao, Zizhen, Hanqing Liu, Fangming Xie, Stephan Fischer, Ricky S. Adkins, Andrew I. Aldridge, Seth A.

Ament, et al. 2021. “A Transcriptomic and Epigenomic Cell Atlas of the Mouse Primary Motor Cortex.” *Nature* 598 (7879): 103–10.
Zhang, Meng, Stephen W. Eichhorn, Brian Zingg, Zizhen Yao, Kaelan Cotter, Hongkui Zeng, Hongwei Dong, and Xiaowei Zhuang. 2021. “Spatially Resolved Cell Atlas of the Mouse Primary Motor Cortex by MERFISH.” *Nature* 598 (7879): 137–43.
Zhang, Yun, Brian D. Aevermann, Trygve E. Bakken, Jeremy A. Miller, Rebecca D. Hodge, Ed S. Lein, and Richard H. Scheuermann. 2021. “FR-Match: Robust Matching of Cell Type Clusters from Single Cell RNA Sequencing Data Using the Friedman-Rafsky Non-Parametric Test.” *Briefings in Bioinformatics* 22 (4). https://doi.org/10.1093/bib/bbaa339.
Zhang, Yun, Brian Aevermann, Rohan Gala, and Richard H. Scheuermann. 2022. “Cell Type Matching in Single-Cell RNA-Sequencing Data Using FR-Match.” *Scientific Reports* 12 (1): 9996.
Zitová, Barbara, and Jan Flusser. 2003. “Image Registration Methods: A Survey.” *Image and Vision Computing*. https://doi.org/10.1016/s0262-8856(03)00137-9.
